# Loss of the s^2^U tRNA modification induces antibiotic tolerance and is linked to changes in ribosomal protein expression

**DOI:** 10.1101/2025.07.01.662621

**Authors:** Katherine L Cotten, Abigail McShane, Jamie C. Smith, Peter C. Dedon, Thomas J. Begley, Kimberly M. Davis

## Abstract

Stress promotes phenotypic changes in bacteria that allow them to survive antibiotic treatment. This phenomenon, termed antibiotic tolerance, can cause treatment failure, highlighting a need to define bacterial pathways that promote survival. Previously, we found *Yersinia pseudotuberculosis* downregulates the gene, *tusB,* in response to doxycycline*. tusB* is involved in generating a sulfur tRNA modification, 2-thiouridine or s^2^U, that increases translation fidelity and efficiency. Here we find that deletion of *tusB* results in loss of s^2^U and induces antibiotic tolerance. Using a combination of sequencing-based approaches and analysis of gene codon usage, our data show that loss of s^2^U decreases translation of ribosomal proteins. Ribosomal proteins are highly enriched in codons that require s^2^U-modified tRNAs for efficient translation, and loss of s^2^U results in ribosome pausing at these codons. Our results highlight a previously unknown mechanism of antibiotic tolerance where reduction in ribosomal protein abundance can globally reduce translation, and describes a novel strategy bacteria use to slow growth by modulating s^2^U levels.

## Introduction

Antibiotic treatment failure is a concerning and multifaceted problem. Major focus has been on antibiotic resistance, which occurs when bacteria acquire a genetic mutation or a genetic element that allows bacteria to grow in the presence of antibiotics^1, 2^. However, bacteria can also develop phenotypic changes in response to stress that allows them to survive antibiotic treatment^1, 2^. This phenomenon is known as antibiotic tolerance when the entire bacterial population responds this way, or antibiotic persistence when only a subpopulation is recalcitrant to antibiotics^3^. These phenotypic changes are transient and typically induce a slowed growth response that renders antibiotics less effective^2^. Once the stress is removed, bacteria resume growth, which is a major concern for relapsing infection^4^. Growing evidence also suggests that antibiotic tolerance and persistence contribute to the development of antibiotic resistance^5–8^. We currently lack strategies to target antibiotic tolerant bacteria, making it important to understand what pathways promote bacterial survival, so we can leverage this information to improve antibiotic efficacy.

*Yersinia pseudotuberculosis* (Yptb) is a Gram-negative bacterium in the class of Gammaproteobacteria that is a natural pathogen of humans and rodents. Yptb has been used as a model to study multiple aspects of pathogenesis of enteric pathogens^9–13^ and recently has been studied in a mouse model of antibiotic persistence^14^. In a previous study we investigated the phenotypic response of Yptb to doxycycline treatment and found expression of the gene *tusB* was significantly downregulated in response to doxycycline^15^. TusB (2-thio<u>u</u>ridine <u>s</u>ubunit <u>B</u>) is involved in a sulfur relay system that transfers a sulfur group onto the wobble position of tRNAs encoding for glutamate (Glu), glutamine (Gln), and lysine (Lys)^16^. TusB complexes with itself and other Tus proteins (TusC and TusD homodimers) forming a heterohexamer that mediates the transfer of sulfur from TusA to TusE^16, 17^. TusE then transfers sulfur to MnmA, which catalyzes the final 2-thiolation reaction, adding a sulfur group onto the second carbon of the uridine in the tRNA anticodon (U34) making 2-thiouridine or s^2^U^16^. This modification restricts binding of the tRNA to purines (A and G), rather than pyrimidines (C or U), thereby increasing translation fidelity and efficiency.

Translation efficiency is generally defined as the rate of protein production per mRNA molecule^18^. Initiation is primarily thought to be the rate limiting step of translation, but elongation can be a limiting step as well^19^. During elongation, the speed at which proteins are translated is impacted by the available pool of tRNAs and competition between cognate and non-cognate tRNAs^20^. tRNA modifications that stabilize the interaction between the anticodon and its cognate codon increase translation efficiency by promoting more rapid decoding. tRNA modifications can thus impact protein translation depending on which codons are decoded by the modified tRNA, and how enriched those codons are in a given transcript^21–23^. In yeast, absence of s^2^U has been shown to impact the metabolic state of cells^24^ and led to proteotoxic stress due to improper protein folding during translation^25^, but has been postulated to not have a major impact on translation efficiency, as measured by correlating transcriptional changes with ribosome footprint changes^24^. In contrast, loss of s^2^U seems to impact translation in *Plasmodium falciparum*, by biasing the translation of certain proteins depending on their enrichment of lysine codons, impacting artemisinin resistance^26^. In *E. coli*, it has been shown that s^2^U deletion mutants have delayed expression of specific proteins, RpoS, Fis, and FtsZ, and impaired cell division^27, 28^. However, it is not known whether s^2^U has a global impact on translation or if this tRNA modification impacts antibiotic susceptibility in bacteria.

Here, we find that deletion of *tusB* and loss of the s^2^U modification renders *Yersinia* tolerant to multiple ribosome-targeting antibiotics, and the RNA polymerase-targeting antibiotic, rifampicin. We hypothesized that loss of the modification slows translation at Glu, Gln, and Lys codons. In line with this, we find that ribosomes pause more frequently at the specific codons that rely on the tRNA modification for proper decoding. We find that ribosomal proteins themselves are highly enriched in these codons and downregulated at the protein level, but not the transcript level, indicating they may be translated less efficiently in the absence of s^2^U. Our results support the model that a reduction in ribosomal proteins is globally reducing translation and is responsible for slowing bacterial growth and inducing antibiotic tolerance.

## Results

### Deletion of *tusB* results in a complete loss of mnm^5^s^2^U and induces changes in other modifications

To understand what impact *tusB* expression has on antibiotic susceptibility, we created a *tusB* deletion strain (Δ*tusB*). First, to confirm that deletion of *tusB* results in loss of the s^2^U modification, we performed chromatography-coupled tandem quadrupole mass spectrometric (LC-MS) analysis to detect relative tRNA modification levels in Δ*tusB* compared to our wild-type (WT) strain (**Figure 1**). We also compared relative tRNA modification levels in WT +/- doxycycline (dox) treatment to see if downregulation of *tusB* would be sufficient to lower tRNA modification levels. We chose to look at tRNA modification levels in the early log phase of growth to match our RNA-seq timepoint where we saw downregulation of *tusB* under WT+dox conditions^15^. Additionally, early to mid-log phase is when the s^2^U modification should be highest in WT cells, prior to nutrient (particularly sulfur) limitation^29^.

**Figure 1.**
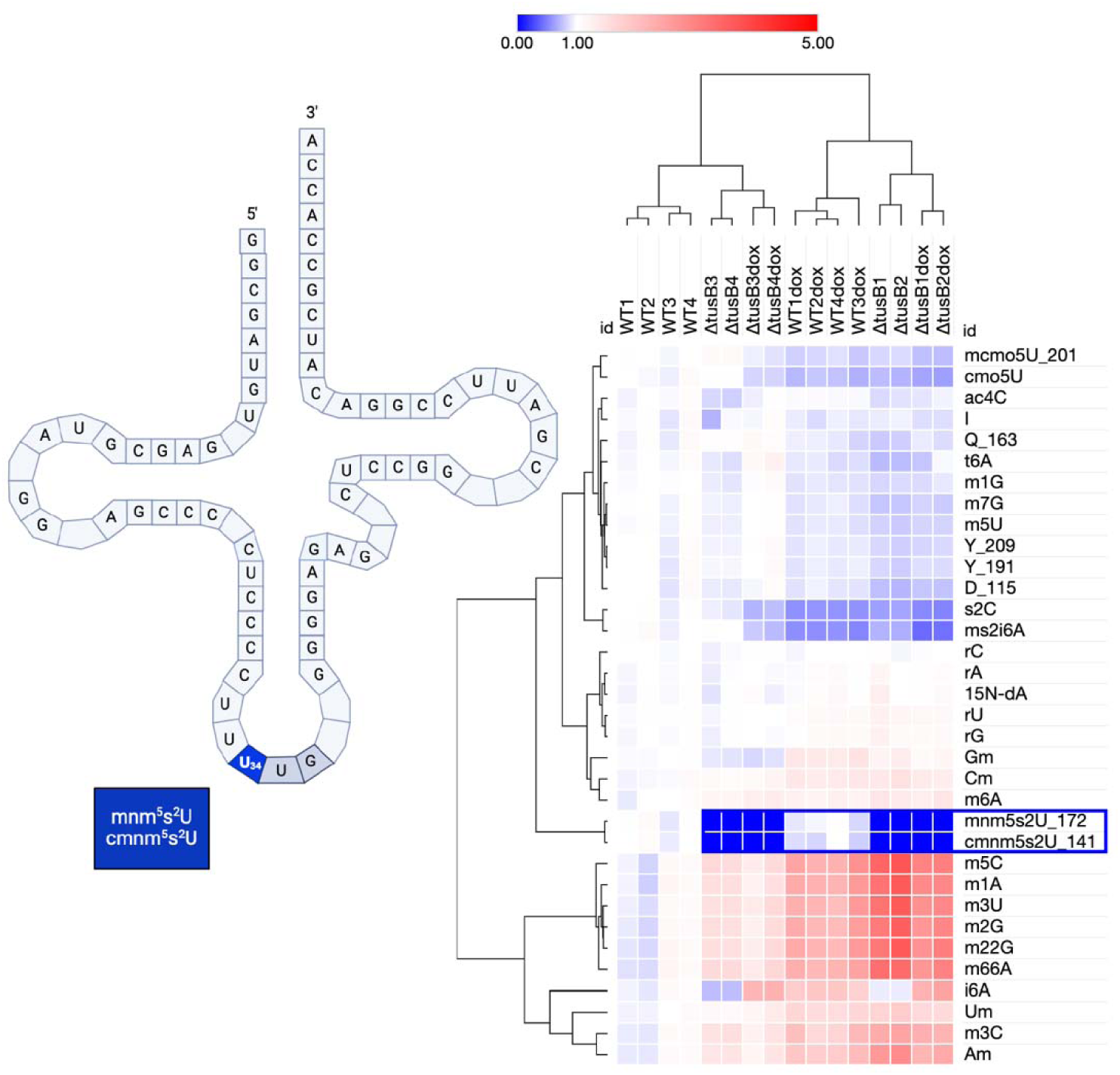
Deletion of *tusB* results in a complete loss of s2U and changes in other tRNA modifications. Log phase bacterial cultures were grown in LB for 2h at 37°C (untreated or treated with 1µg/mL of doxycycline). The *tusB* deletion strain (Δ*tusB*) grows 2-fold slower than the WT strain. Δ*tusB* cultures were scaled up so all samples had approximately 10^9^ CFUs. Small RNA was purified and digested into nucleosides. Changes in relative quantities of modified nucleosides were measured by LC-MS/MS. Each sample is normalized to the average peak area of four biological replicates for WT untreated (WT1-4), see Supplemental Data 1. Left: Created in BioRender. Cotten, K. (2027) https://BioRender.com/kyv58wc

Hierarchal clustering analysis shows that WT samples cluster separately based on treatment. Oddly *tusB* deletion strains cluster together regardless of treatment, which may in part be due to batch effect (**Figure 1**). Although specific Dox-dependent differences were also detected and are highlighted below (i6A). Importantly, as expected, we found a complete loss of s^2^U in Δ*tusB* (**Figure 1, blue box**). We saw minimal downregulation of 5-methylaminomethyl-2-thiouridine (mnm^5^s^2^U, average 0.9-fold change WT dox vs. WT untreated) and 5-carboxymethylaminomethyl-2-thiouridine (cmnm^5^s^2^U, average 0.87-fold change WT dox vs. WT untreated) in response to doxycycline.

Interestingly we detected other changes in relative tRNA modification levels in Δ*tusB* compared to WT, and in response to doxycycline treatment. Two sulfur tRNA modifications, 2-thiocytidine (s^2^C) and 2-methylthio-N6-isopentenyladenosine (ms^2^i6A), were downregulated in response to doxycycline (both in WT and Δ*tusB*) and somewhat in response to *tusB* deletion alone (**Figure 1**), which could indicate low sulfur availability^29^. Additionally, we also detected upregulation of many modifications commonly found on ribosomal RNA (m^5^C, m^1^A, m^3^U, m^2^G, and m^66^A). Our RNA isolation enriches for tRNA based on size and can include smaller ribosomal RNA fragments if these are abundant.

### Loss of 2-thiolation tRNA modification induces tolerance to specific ribosome and RNA-polymerase targeting antibiotics

We tested the growth and antibiotic susceptibility of Δ*tusB* and found that it was significantly slower growing and more tolerant to multiple antibiotics including the ribosome-targeting antibiotics, doxycycline and gentamicin, and the RNA polymerase-targeting antibiotic, rifampicin (**Figure 2**). We detected a slight decrease in antibiotic susceptibility to chloramphenicol, although this was not statistically significant. These phenotypes were rescued by integrating the *tusB* gene back into the genome (*tusB* rescue, **Figure 2**). Additionally, to confirm that phenotypes were specific to loss of the tRNA s^2^U modification, we complemented the deletion strain with the *Bacillus subtilis* pathway for 2-thiolation. *B. subtilis* possess an abbreviated version of the sulfur relay system where the cysteine desulfurase, YrvO, transfers sulfur groups directly to MnmA, which catalyzes the final 2-thiolation reaction^30^. As expected, expression of *yrvO-mnmA* restored growth and antibiotic susceptibility of Δ*tusB*, although this was not statistically significant for doxycycline (**Figure 2**). Chloramphenicol susceptibility was similar to Δ*tusB* when *yrvO-mnmA* was expressed, indicating that loss of the modification does not greatly impact chloramphenicol susceptibility. Importantly, phenotypes were only complemented when a functional copy of *yrvO* was expressed and not when the active cysteine residue (C325A) of YrvO was mutated or with an empty vector control (**Extended Data Figure 2**)^30^.

**Figure 2.**
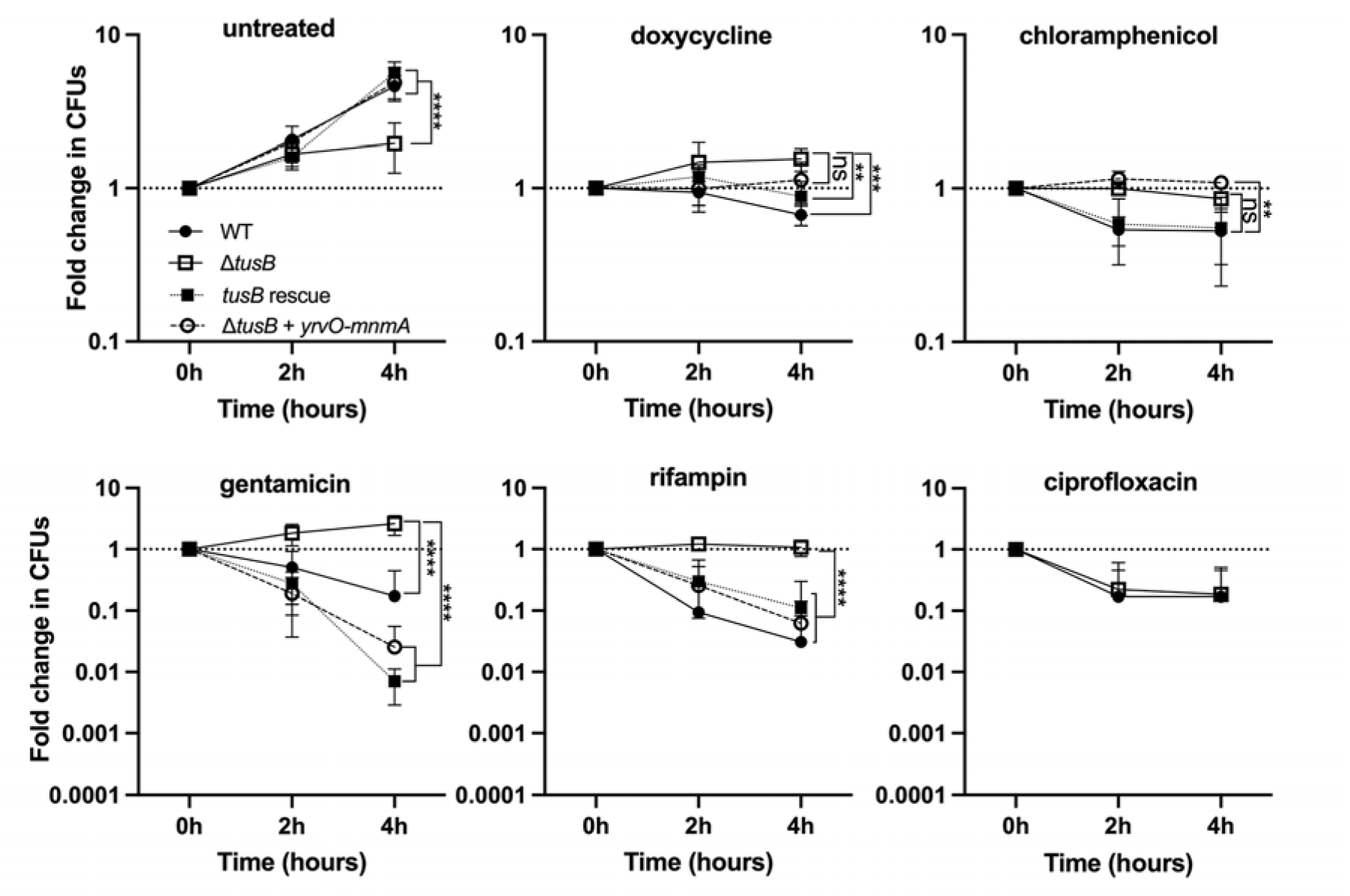
Deletion of *tusB* induces slowed growth and antibiotic tolerance to ribosome and RNA-polymerase targeting antibiotics due to loss of s^2^U tRNA modification. Log phase bacterial cultures were grown at 37C and sampled every 2h to enumerate CFUs (colony forming units). *tusB* rescue was generated by introducing the *tusB* gene back into its native site in the genome by homologous recombination. Complemented strains carry a plasmid (pDS145)^30^ with the *yrvO-mnmA* open reading frames under the control of the P*_ara_*promoter, and were treated with 0.2% arabinose throughout the experiment to induce expression of the construct. Cultures were treated with either 1µg/mL doxycycline, 25µg/mL chloramphenicol, 2µg/mL gentamicin, 2µg/mL rifampicin, 0.1µg/mL ciprofloxacin, or left untreated. Fold change in CFUs are shown by normalizing each timepoint to the 0h timepoint. Two-way ANOVA with Tukey’s multiple comparisons. Asterisks denote significant p-values as follows: **** p <0.0001, *** p <0.001, ** p < 0.01, and * p < 0.050. Data points represent the mean of three biological replicates, error bars represent standard deviation (SD). See Supplemental Data 2.

To understand if overexpressing one of the affected tRNAs could overcompensate the lack of s^2^U we complemented the *tusB* deletion strain with a plasmid overexpressing the lysine (UUU) tRNA gene. Overexpression of lysine (UUU) tRNA was able to compensate for the lack of s^2^U in yeast, restoring growth^31^. However, this had no effect on growth or antibiotic susceptibility in the *tusB* deletion strain (**Extended Data Figure 3**), likely because it did not account for the effect of loss of s^2^U on the other tRNAs encoding for glutamine and glutamate.

Overall, these results indicate that loss of the s^2^U tRNA modification induces tolerance to the ribosome and RNA-polymerase targeting antibiotics that were tested. Interestingly, we observed no difference in susceptibility of Δ*tusB* to the DNA-gyrase targeting antibiotic, ciprofloxacin, (**Figure 2**) indicating that this tolerance phenotype may be specific to antibiotics that target translation and transcription. Since transcription and translation are generally coupled in bacteria, we hypothesized that this might indicate translation is impacted by loss of s^2^U.

### Δ*tusB* has a starvation-like metabolic signature and decreased levels of ribosomal proteins

To further understand the impact of loss of s^2^U, we performed RNA-seq and mass spectrometry to define the transcriptome and proteome, respectively, of Δ*tusB* compared to WT. 259 genes were significantly downregulated, and 182 genes were significantly upregulated at the transcript level (p_adj_ < 0.05 and >2-fold change). At the protein level, 166 genes were significantly downregulated, and 197 genes were significantly upregulated (p-value < 0.05 and >2-fold change in protein abundance).

KEGG Pathway Enrichment analysis of differentially regulated genes from our RNA-seq dataset shows that Δ*tusB* upregulates multiple amino acid biosynthesis pathways (**Figure 3A**). Many of these are also upregulated at the protein level in our mass spectrometry dataset (**Figure 3B**). We hypothesized this could indicate the stringent response is being induced by loss of the tRNA modification. *In vitro* studies have shown the s^2^U modification improves tRNA synthetase binding^32^ making it plausible that loss of the modification could result in uncharged tRNAs binding to the ribosome, inducing the stringent response. However, deleting the alarmone synthetase, *relA*, in the *tusB* deletion background did not rescue the slowed growth or antibiotic tolerance phenotype (**Extended Data Figure 4**), indicating the Δ*tusB* phenotype is not linked to the stringent response. We also saw no indications that the SOS response, which is another stress response pathway known to induce antibiotic tolerance (via inhibition of cell division in response to DNA damage) was upregulated in *tusB* deletion. For example, transcript and protein levels of key regulators of the SOS response including *sulA*, (which was not detected by RNA-seq or mass spec) and *recA* (which was slightly downregulated by RNA-seq, -0.4 log2fold change, and was not significantly different by mass spec between WT and Δ*tusB*) indicated that the SOS response is likely not induced (see Supplemental Data 3).

**Figure 3.**
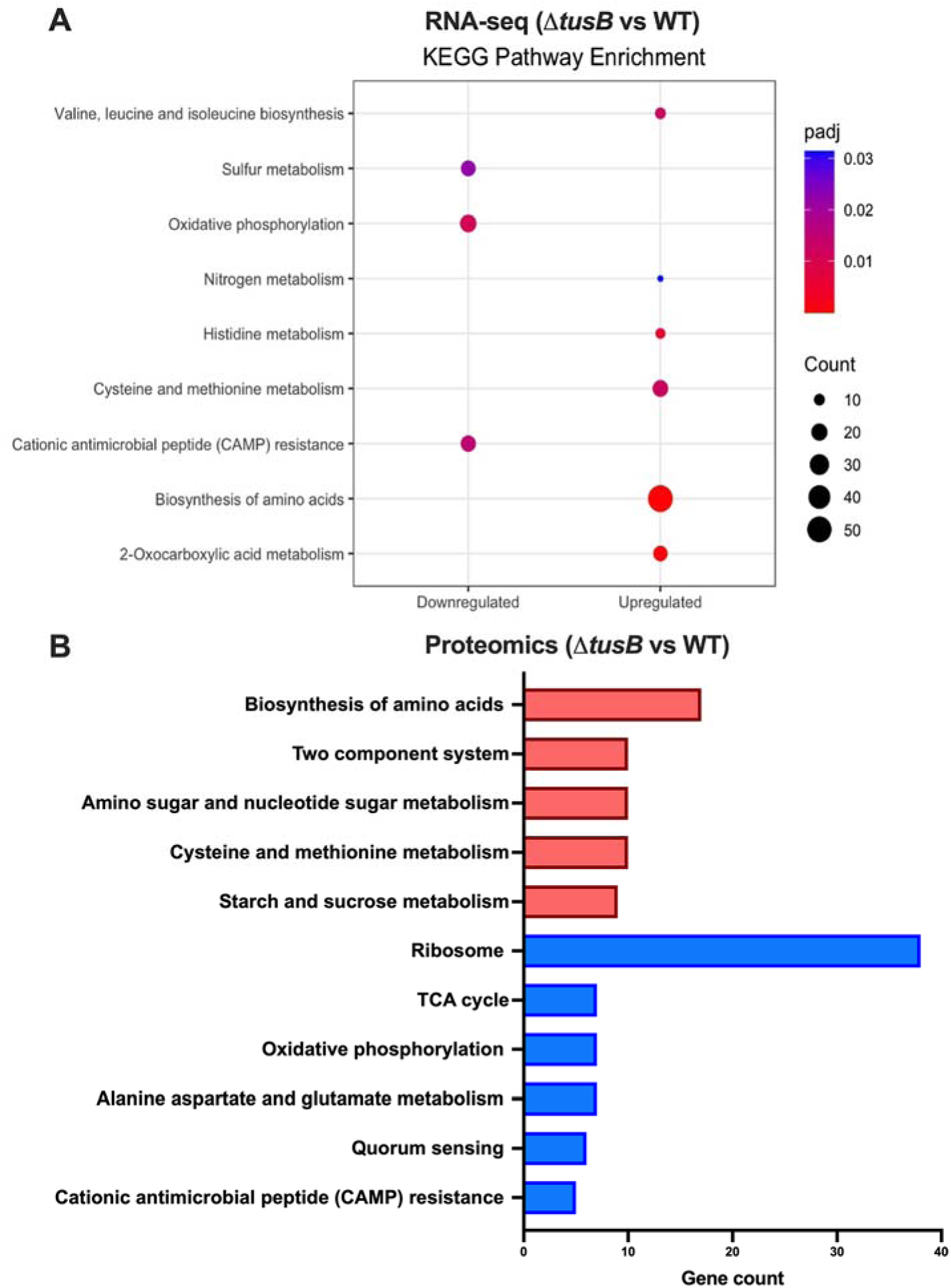
Δ*tusB* has a starvation like metabolic profile and reduced ribosomal protein levels. **A.** Log phase bacterial cultures were grown for 2h at 37°C (approximately 10^8^ CFUs) before isolating RNA. Significant changes in transcript levels between WT and Δ*tusB* were evaluated by DESeq2. Enriched KEGG Pathways (p-value < 0.05) are shown for downregulated and upregulated genes. The number of genes associated with each pathway is represented by circle size. Three biological replicates. **B.** Protein was isolated from bacterial cultures after 2h of growth at 37°C (approximately 10^8^ CFUs). Significant changes in protein abundance between WT and Δ*tusB* were evaluated by One-way ANOVA. The number of significantly regulated genes (p-value < 0.05) associated with KEGG pathways are shown. Three biological replicates. See Supplemental Data 3.

RNA-seq and proteomic analyses generally revealed that Δ*tusB* has a starvation-like metabolic signature. Δ*tusB* significantly downregulated oxidative phosphorylation (**Figure 3A, B)** as indicated by downregulation of several genes in the TCA cycle and multiple subunits of the electron transport chain. This type of metabolic shift is often associated with antibiotic tolerance and persistence and is consistent with the slowed growth phenotype of Δ*tusB*.

Surprisingly, our proteomic analysis revealed that 37 of the 54 ribosomal protein subunits were significantly downregulated in Δ*tusB* (**Figure 3B**). Most of these ribosomal subunits did not show up as hits in our RNA-seq dataset and the few that did (p<0.05) were not strongly downregulated or upregulated at the transcript level (less than 2-fold change) (**Extended Data Table 1**). This also indicates that the stringent response is likely not be responsible for the expression changes in Δ*tusB*, since the ppGpp alarmone has been shown to repress transcription of ribosomal protein genes^33^. We hypothesized that this difference between our RNA-seq and proteomics datasets could indicate that the ability to translate these proteins is impaired in Δ*tusB*.

### Ribosomes pause more frequently at Glu, Gln, and Lys codons in Δ*tusB*

To understand what impact *tusB* deletion has on translation, we performed ribosome profiling to determine if loss of the s^2^U tRNA modification impacts ribosome speed. We adopted a ribosome profiling approach developed by Mohammad et al.^34, 35^, allowing us to identify ribosome abundance and positioning at single codon resolution. Ribosome pause scores are calculated by taking the average ribosome density at a single codon normalized to the average ribosome density across a given gene and averaging this for all genes in the genome. As expected, the relative ribosome codon occupancy (Δ*tusB* pause score normalized to WT) was moderately increased at Glu, (GAG and GAA), Gln, (CAA), and Lys (AAA and AAG), indicating that ribosomes paused more frequently at these codons in Δ*tusB* (**Figure 4**). No change in ribosome pausing at CAG was expected since Yptb have one Gln tRNA that is not modified with s^2^U (tRNA^CUG^) that is likely sufficient for decoding CAG^36^. In contrast, Yptb seem to more heavily rely on modified tRNAs for decoding Lys (4 tRNA^UUU^ and only one tRNA^CUU^)^36^ and Glu (5 tRNA^UUC^ and no tRNA^CUC^)^36^, which explains why we see ribosome pausing at both isoacceptor codons for Lys and Glu.

**Figure 4.**
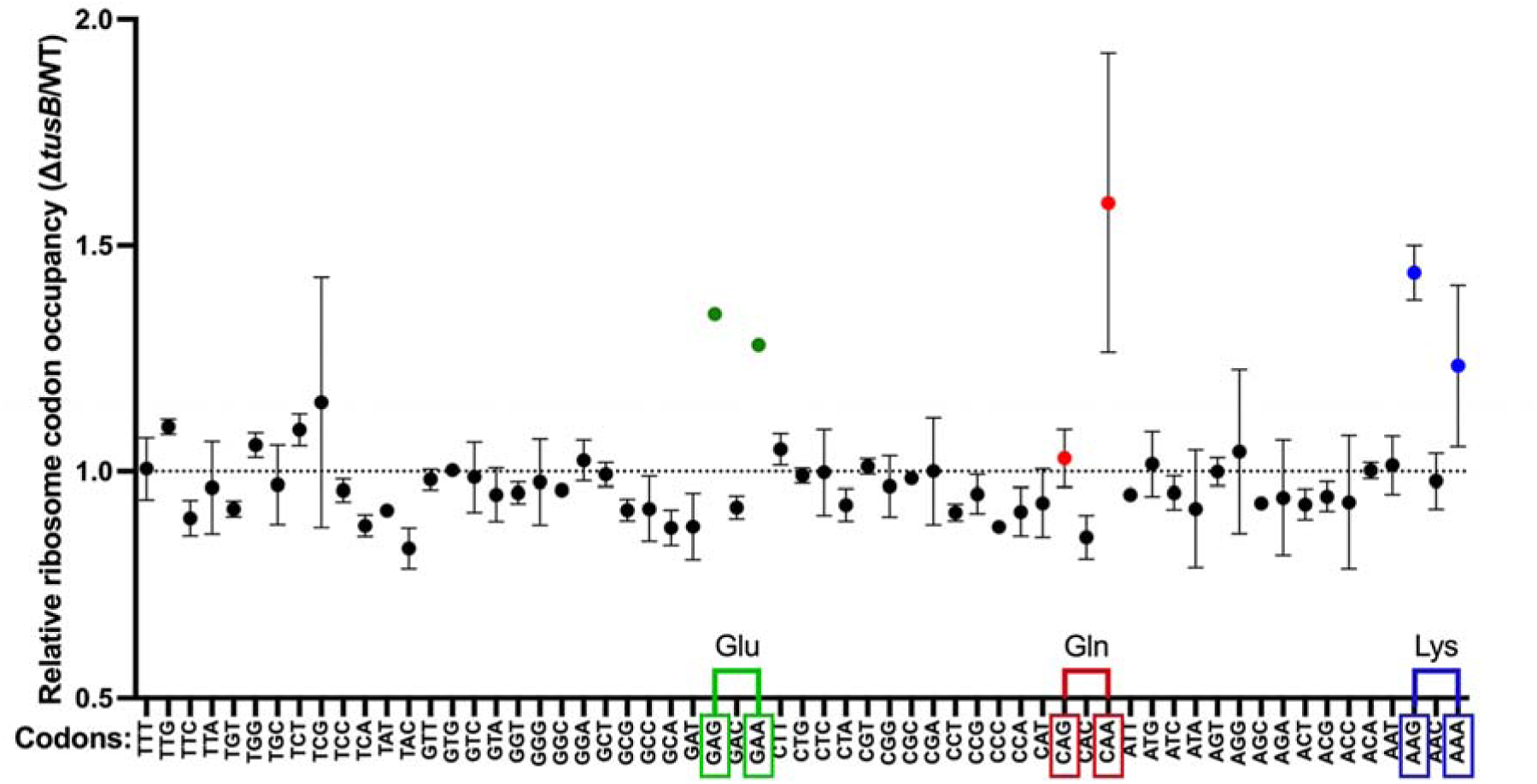
Ribosomes pause more frequently at Glutamate, Glutamine, and Lysine codons. Cultures of bacteria were grown for 5h at 37°C (approximately 10^10^ CFUs) and then flash frozen in liquid nitrogen to stop translation. Ribosome protected fragments were isolated and sequenced to measure the abundance of ribosomes at each codon. Pause scores are calculated by taking the average ribosome density at a codon normalized to the average ribosome density across a given gene for all genes in the genome. Relative ribosome occupancy (pause score of Δ*tusB* normalized to the pause score of WT) is shown for each codon. Data points represent the mean of two biological replicates, error bars represent standard deviation (SD). See Supplemental Data 4.

In addition to indicating slower translation, ribosome pausing can also be a sign that proteins are being mistranslated and/or not properly folded^25^. Since s^2^U is important for stabilizing the interaction between the tRNA anticodon and its cognate codon, we briefly investigated if the absence of s^2^U impacts translation fidelity, with the idea that loss of s^2^U would increase the chances that a near cognate tRNA could bind and incorporate the wrong amino acid. We detected potential misincorporation of amino acids in a preliminary LC-MS/MS run, considering the following changes as modifications in peptides: glutamate to aspartate, glutamine to histidine, and lysine to asparagine. To understand if these misincorporations occur more frequently due to loss of s^2^U, we measured the relative abundance of modified to unmodified peptides in Δ*tusB* vs. WT (**Extended Data Figure 5A**, **Supplemental Data 8**). Interestingly we detected on average higher frequency of misincorporation at Glutamine CAA versus Glutamine CAG codons in Δ*tusB* compared to WT (**Extended Data Figure 5B**), consistent with our finding that ribosomes pause at CAA and not CAG (**Figure 4**). However, with a small sample size we cannot conclusively say whether misincorporation of amino acids is more frequent due to loss of s^2^U. Additionally, it does not seem that proteins are being widely mistranslated in Δ*tusB*, given that we saw no indication of proteotoxic stress in our differential expression analysis (**Figure 3A, B**).

Overall, our results indicate that absence of s^2^U impacts ribosome speed. The ribosome pausing we detected at Glu, Gln, and Lys codons (**Figure 4**) is comparable to what has been reported in yeast^25, 37^, although we detected relatively higher pause scores at Gln CAA and Lys AAG codons. These values are considered a moderate increase in ribosome occupancy, indicating that the ribosome is transiently pausing at these sites rather than stalling. Although transient pausing at one site may not have a great impact on protein output, we hypothesized that if transcripts were highly enriched in these codons, this would significantly slow down the ribosome and decrease translation of a given transcript.

### Switching Gln codons from CAA to CAG slightly improves translation of GFP

To test whether enrichment of codons with ribosome pausing is sufficient to slow translation and impact protein output, we created a codon-optimized GFP reporter construct where all CAA codons (6/8 total Gln codons) are switched to CAG (**Figure 5A**). We focused on Gln codons since we saw a clear increase in ribosome pausing at CAA, but not CAG (**Figure 4**). As expected, Δ*tusB* grew slower than WT (**Figure 5B**). However, Δ*tusB* growth was slightly improved when expressing the CAG construct (**Figure 5B**), consistent with others have reports that bacterial growth is impaired by expressing *gfp* with codons that slow the ribosome^20^. Oddly, we saw lower GFP fluorescence from the CAG construct in the WT strain compared to native GFP, which was unexpected since CAA and CAG are used equally in protein coding sequences in Yptb and thus switching these codons was not predicted to bias translation in the WT strain (**Figure 5C**). However, GFP fluorescence from the CAG construct was not lower than native GFP in Δ*tusB*, instead it was slightly higher (**Figure 5C**). As a proxy for GFP translation, we quantified GFP fluorescence relative to *gfp* transcript levels at 4h (**Figure 5D**). Importantly we can see GFP fluorescence per relative *gfp* transcript level is trending lower in Δ*tusB* vs. WT (p-value = 0.0552) for native GFP, suggesting GFP translation is lower in Δ*tusB*. GFP fluorescence per relative *gfp* transcript level was slightly improved in Δ*tusB* with the CAG construct, suggesting that replacing CAA codons with CAG slightly improves GFP translation, although this was not statistically significant.

**Figure 5.**
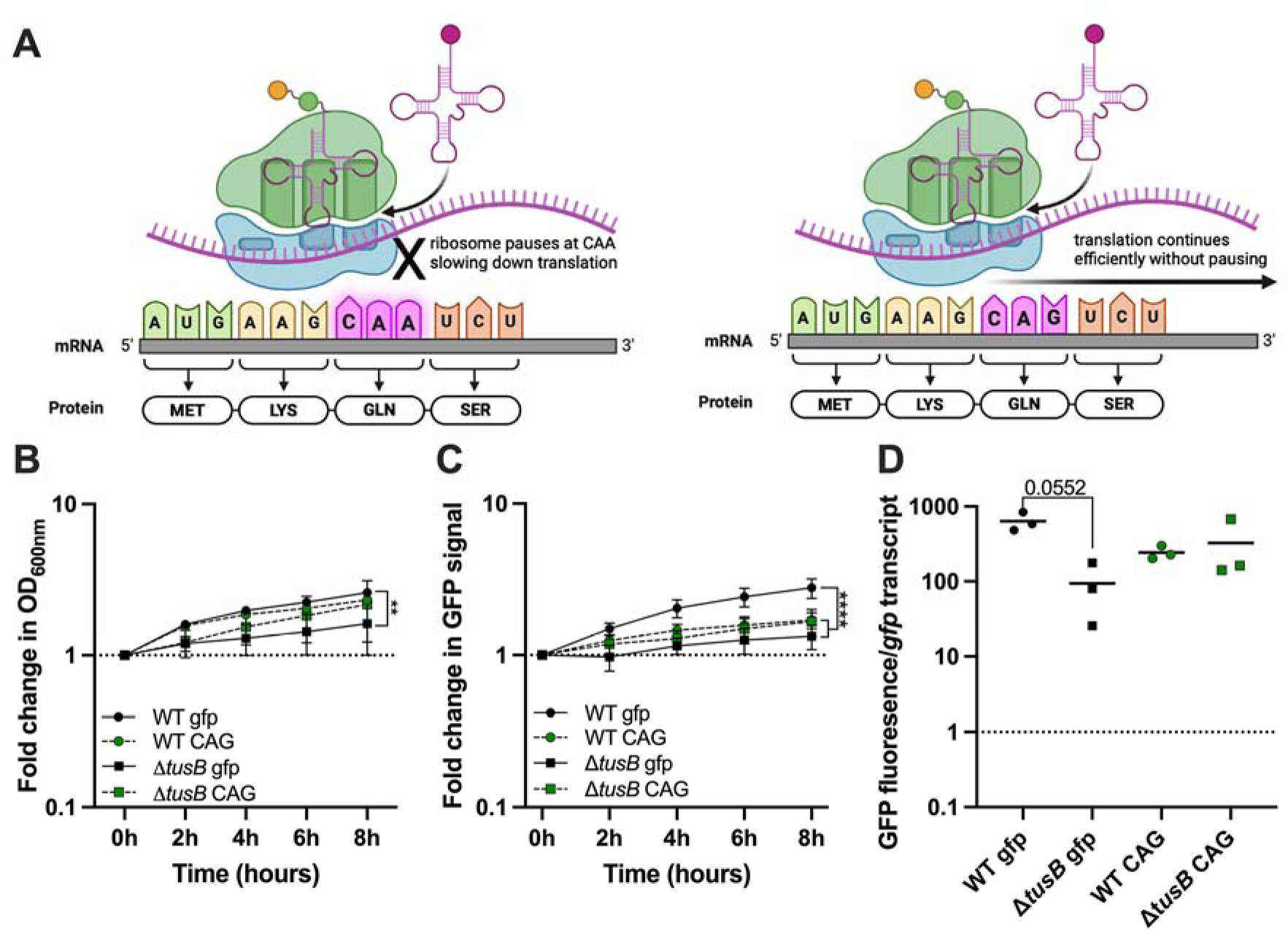
Switching Glutamine codons from CAA to CAG slightly improves GFP translation. **A.** Graphical depiction of model. **B, C, D.** Log phase bacterial cultures were grown for 8h at 37°C. At 0h cultures were treated with 1mM IPTG to induce expression of *gfp* or left untreated as a control. Created in BioRender. Cotten, K. (2026) https://BioRender.com/z1zzgwq **B.** Fold change in OD600 is shown by normalizing each timepoint to the 0h timepoint. **C.** GFP fluorescence is normalized to optical density (OD600) to account for differences in bacterial density between samples. Values are normalized to the 0h timepoint to show fold change in GFP fluorescence overtime from the point of IPTG induction. **D.** RNA was isolated at 4h during the experiment and *gfp* transcript levels were quantified by qPCR. GFP fluorescence is normalized to *gfp* transcript level as a proxy for GFP protein translation. GFP fluorescence is normalized to the untreated (uninduced) control to account for background autofluorescence of the bacteria. *gfp* transcript levels were also normalized to the untreated (uninduced) control to account for any leaky expression of the construct. For B and C, two-way ANOVA with Tukey’s multiple comparisons. Asterisks denote significant p-values as follows: **** p <0.0001, ** p < 0.01. Data points represent the mean of three biological replicates, error bars represent standard deviation (SD). For D, one-way ANOVA with Kruskal-Wallis test for multiple comparisons. Line represents the mean of three biological replicates. See Supplemental Data 5.

Overall, these results indicate that translation is lower when s^2^U is absent and is partially improved by replacing CAA codons with CAG. We expect that GFP translation was not significantly improved by switching only Glutamine codons since the other codons, Glutamate and Lysine, likely also contribute to the lower translation of GFP in Δ*tusB*

Since GFP protein levels can also be impacted by protein decay and degradation, we measured GFP decay using a ssrA tag to target the GFP protein to the ClpXP proteosome^38, 39^. Unstable GFP (*gfp-ssrA*) had a half-life of 57 minutes in WT. In contrast, unstable GFP (*gfp-ssrA*) had a half-life of 187 minutes in Δ*tusB*, indicating that protein degradation is impaired (**Extended Data Figure 6**). Additionally, stable GFP also had a longer half-life in Δ*tusB* (277 minutes) compared to WT (165 minutes) (**Extended Data Figure 6**). These findings may be explained by our RNA-seq and proteomic analysis which indicates that Δ*tusB* downregulates the ClpXP proteosome (*clpX* is downregulated 1.79-fold by RNA-seq (padj =1×10^8^) and 3-fold in protein abundance (p = 0.0292); *clpP* is downregulated 1.69-fold by RNA-seq (padj = 1.49×10^8^) and 3.77-fold in protein abundance (p=0.00508), see **Supplemental Data 9**). It is unclear why Δ*tusB* downregulates the ClpXP proteasome, but this suggests that Δ*tusB* is impaired in protein degradation, allowing us to conclude that the lower GFP fluorescence of Δ*tusB* (**Figure 5C, D**) is not due to increased degradation of GFP proteins.

### Lys AAG codon enrichment is highly associated with subset of downregulated ribosomal proteins

To look globally at the impact of loss of the s^2^U tRNA modification on biasing translation, we analyzed codon usage for all protein-coding genes in the *Yersinia pseudotuberculosis* IP2666 genome, with the idea that enrichment of Glu, Gln, and Lys codons would be highly associated with downregulated proteins if these codons were biasing translation. Codon Z-scores, detailing relative codon over-or under-use in a gene (relative to genome averages), were calculated for each codon in each gene by subtracting the average global codon frequency from the gene codon frequency and normalizing to the standard deviation of that codon’s usage throughout the genome^40^.

As a result, codon Z-scores represent how many standard deviations above or below the global frequency a specific codon is used in a given gene. We performed principal component (PC) analysis using the codon Z-scores for all genes that were differentially regulated at the protein level (2-fold change in protein abundance, Δ*tusB* vs. WT, p-value < 0.05). PC scores are calculated for each gene based on their codon usage and plotted in **Figure 6A**, revealing a cluster of downregulated proteins along PC1. We performed principal component regression analysis using the log2fold change in protein abundance (Δ*tusB* vs. WT) as the dependent variable and found several codons that significantly contribute to the variance in protein expression. Codons of interest, Glu GAA (p-value=0.0345), Lys AAG (p-value <0.0001) and Lys AAA (although not significant, p-value = 0.1757) corresponded with the cluster of downregulated proteins along PC1 in the lower right quadrant of the loadings plot (**Figure 6B, C**). The codon Z-scores are particularly high for Lys AAG amongst the downregulated proteins in this subset. Interestingly, 18 of the 21 downregulated proteins are ribosomal proteins, indicating that enrichment of Lys AAG codons may explain why these proteins are less abundant even though none were significantly downregulated at the transcript level (**Figure 3B**). Although we were also expecting Gln CAA codons to bias translation of proteins, principal component regression analysis indicated they did not significantly contribute to the variance.

**Figure 6.**
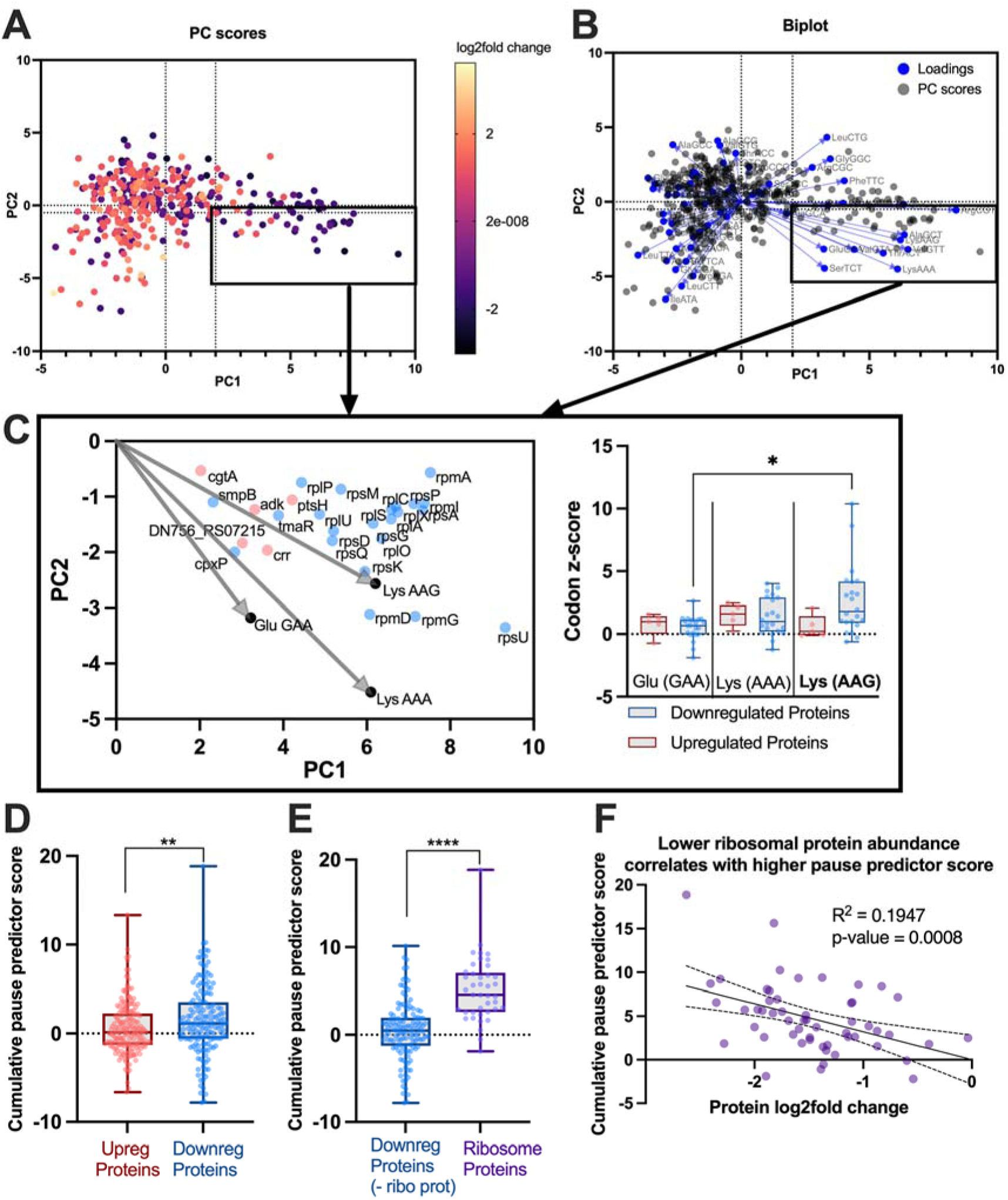
A combination of Glu, Gln, and Lys codons are highly associated with ribosomal proteins. Codon Z-scores were calculated for each gene by taking the frequency of a given codon in that gene and normalizing it the global frequency of that codon and the standard deviation of that codon. **A.** A data matrix of codon Z-scores for all genes that were significantly regulated at the protein level in Δ*tusB* vs. WT (2-fold change, p<0.05) were analyzed by principal component analysis. PC scores are plotted for each gene based on their codon usage. PC1 explains 11.3% of the variance and PC2 explains 6.4% of the variance. **B.** Biplot shows the loadings plot, linear association of each codon, with the PC scores of each gene. **C.** Biplot of genes associated with Lys AAG, Lys AAA, and Glu GAA (PC score >2 and <- 0.5) are graphed separately for clarity. Boxplots of codon Z-scores amongst upregulated (red) and downregulated (blue) genes in this subset are shown for Glu GAA, Lys AAA and Lys AAG. One-way ANOVA with Dunnett’s T3 multiple comparisons (* denotes p < 0.05). **D.** Cumulative pause predictor scores were calculated by taking the sum of Glu (GAA), Glu (GAG), Gln (CAA), Lys (AAA), Lys (AAG), codon Z-scores weighted by their ribosome pause score. Cumulative pause predictor scores are higher on average amongst downregulated proteins (blue) versus upregulated proteins (red). Mann Whitney test, ** p < 0.01. **E.** Ribosomal proteins that are significantly downregulated at the protein level (purple) have on average higher cumulative pause predictor scores compared to all other downregulated proteins (blue). Mann Whitney test, *** p < 0.0001. **F.** The log2fold change in protein abundance in Δ*tusB* vs. WT (x-axis) is plotted against the cumulative pause predictor score for all 54 ribosomal proteins. Simple linear regression shows a negative slope (−3.215) and significant deviation from zero (p-value = 0.0008). See Supplemental Data 6.

### A combination of Glu, Gln and Lys enrichment likely contributes to decreased ribosomal protein levels

Finally, we also looked at the combined impact of all five codons that increased ribosome pausing, with the idea that enrichment of a combination of Glu, Gln (CAA) and Lys codons would have more impact than enrichment of a single codon, and that codons with higher ribosome pause scores would have a greater impact on the translation of a given protein. We took the sum of the codon Z-scores for Glu (GAA), Glu (GAG), Gln (CAA), Lys (AAA), and Lys (AAG) for each gene and weighted (multiplied) each codon Z-score by the average ribosome pause score for that respective codon. We refer to this calculation as the “cumulative pause predictor score”, with the idea that a higher score will predict a gene will be translated less efficiently due to increased ribosome pausing along the transcript. Consistent with this idea, we saw significantly increased cumulative pause predictor scores among proteins that were downregulated in Δ*tusB* compared to upregulated proteins (**Figure 6D**). Ribosomal proteins in particular had significantly higher cumulative pause predictor scores than all other downregulated proteins (**Figure 6E**), suggesting that not only enrichment of Lys AAG, but likely enrichment of a combination of all five of these codons is slowing down the translation of ribosomal proteins when s^2^U is absent. Importantly, we do not see this same trend in five other codons selected as a control that did not have changes in ribosome pausing (**Extended Data Figure 7**). Additionally, for the 54 ribosomal proteins, we see that higher cumulative pause predictor scores correlate with decreased protein abundance (**Figure 6F**).

Overall, these results suggest that translation of ribosomal proteins is significantly reduced when s^2^U is lost due to an enrichment of codons that slow down the ribosome.

### Loss of s2U results in slow growth and antibiotic tolerance *in vivo*

Since broth media conditions do not mimic the complex environment of a host during infection, we aimed to test the growth and antibiotic susceptibility of the *tusB* deletion strain *in vivo* using a mouse model of infection. In previous studies^14,15^ we have utilized a systemic model of infection, wherein C57BL/6 mice are inoculated with 10^3^ CFUs via tail vein injection (Figure 7A). Bacteria spread systemically to sites such as the spleen and after 48 hours distinct microcolonies are formed, which are centers of bacteria surrounded by circumscribing layers of neutrophils and macrophages^10, 14^. At 48 hours an average bacterial load of 10^5^ CFUs is reached and by 72 hours It reaches 10^6^ CFUs. Doxycycline treatment at the 48-hour timepoint results in a 10-fold drop in CFUs^14^. In contrast, the *tusB* deletion strain is severely attenuated in the mouse model, only reaching 10^2^ CFUs (Figure 7B). Due to this severe attenuation, 10x higher dose was used to infect mice. With this dose bacterial load was 10x higher, but still only reached ∼10^3^ CFUs 48 hours post infection (Figure 7C). As expected, bacteria were slow growing resulting in no change in bacterial load between 48 and 72 hours post infection, and bacteria were tolerant to doxycycline, resulting in no drop in bacterial load after doxycycline treatment (Figure 7C).

**Figure 7.**
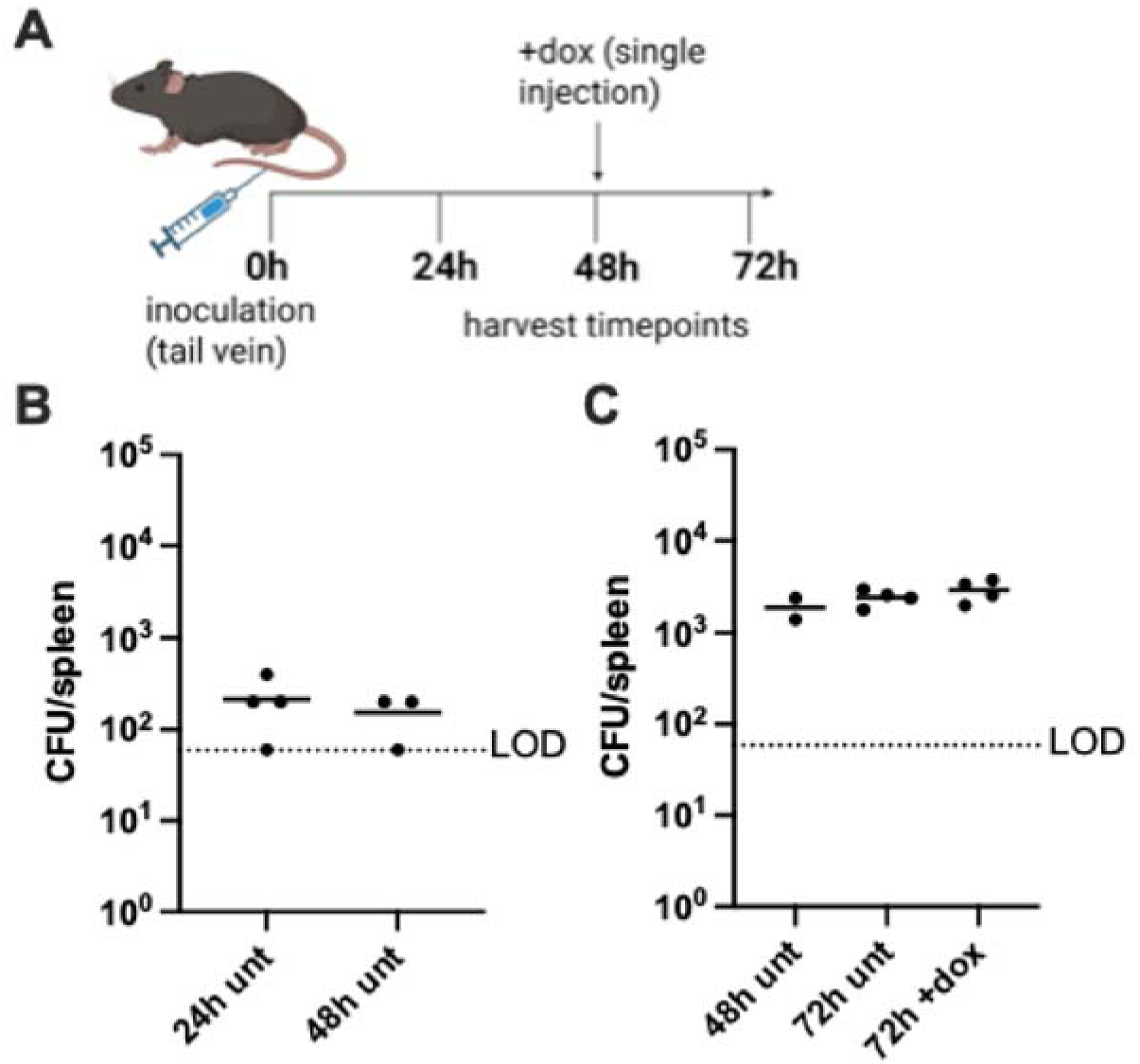
Deletion of *tusB* induces slowed growth and antibiotic tolerance in a mouse model of infection. (A) Graphical timeline of the mouse infection. C57BL/6 mice were inoculated with either 10^3^ or 10^4^ CFUs by tail vein injection. After 48h post infection mice were either treated with 40kg/mg of doxycycline by intraperitoneal injection or left untreated. Spleens were harvested at the indicated timepoints to enumerate CFUs. Created in BioRender. Cotten, K. (2027) https://BioRender.com/b41q462 (B) CFU quantification is shown from mice infected with 10^3^ CFUs. (C) CFU quantification is shown from mice infected with 10^4^ CFUs. **See Supplemental Data.**

## Discussion

This work provides evidence for a novel mechanism by which loss of s^2^U induces antibiotic tolerance. It also adds to the growing evidence that modulation of tRNA modifications can widely impact expression of genes through codon biases in translation^21, 23, 41^. In other reports, s^2^U has been assumed to not have a drastic impact on protein translation output when only a moderate increase in ribosome pausing is seen^18,19,25^. However, here we see ribosome pausing highly correlates with genes that are significantly downregulated at the protein level, but not at the transcript level.

Although pausing at one site alone is likely not sufficient to significantly slow down translation, our results suggest that enrichment of ribosome pausing sites can lead to reduced protein levels. Ribosomal proteins seem to be the most impacted by this codon bias in translation as indicated by their high enrichment in Glu, Gln, and Lys codons and the significant reduction in ribosomal protein levels. We expect that this reduction in ribosomal proteins is globally reducing the translational capacity of the cell and is responsible for inducing tolerance to ribosome and RNA polymerase-targeting antibiotics. However, how exactly slowed translation improves antibiotic survival remains an open question.

The s^2^U tRNA modification is well conserved among bacteria^16^ and ribosomal protein sequences are also generally conserved across bacterial species, including prominent ESKAPE pathogens that are highly resistant to antibiotics (**Extended Data Figure 8**). Therefore, it is plausible that this could be an antibiotic tolerance mechanism used broadly by bacteria to better survive antibiotics, although this would depend on the codon usage and tRNA availability for each species.

Future work will be needed to understand what conditions in the host environment are sufficient to reduce s^2^U levels. In our study doxycycline treatment only moderately lowered s^2^U tRNA modification levels (**Figure 1**). However, given the broad downregulation of the 2-thiolation pathway in response to gentamicin (**Extended Data Figure 1**), we expect that gentamicin may have a stronger impact on modulating s^2^U levels and promoting survival.

Given that s^2^U levels are modulated in response to sulfur availability^29^, it is likely that conditions within the host environment, other than antibiotic stress, also result in lowered s^2^U levels within bacterial pathogens. This highlights that the mechanism we have defined here may be one used broadly by bacteria to adapt to conditions where it is beneficial to slow growth, such as low nutrient conditions. Fast growing cells need to have a high number of ribosomes since protein synthesis is limited by the availability of free ribosomes ^42^, and ribosome abundance is linearly correlated with growth rate^43^.

However, during slow growth, it is beneficial for bacteria to slow ribosome protein synthesis since ribosome biogenesis and translation are energetically demanding processes. Therefore, it is interesting to speculate that the impact of s^2^U on ribosomal protein levels we see here could be a previously unknown mechanism by which bacteria reduce translation of ribosomal proteins in response to conditions where it is beneficial to slow growth and save energy.

Surprisingly we detected some changes in other tRNA modifications (**Figure 1**), some of which are modifications also found on ribosomal RNA (rRNA). Particularly under starvation conditions, ribosomal RNA is more highly degraded, which could explain why we detected increased levels of modifications commonly found on rRNA in Δ*tusB* ^40,41^. However, it is unlikely that rRNA degradation is responsible for lower ribosomal protein levels in Δ*tusB*. 16S rRNA levels were not different by qPCR in WT versus Δ*tusB* (**see Supplemental Data 7**). Additionally, given the reduced capability of Δ*tusB* to degrade GFP (**Extended Data Figure 6**), along with downregulation of the ClpXP protease, it is unlikely that ribosomal proteins are being readily degraded in Δ*tusB*. Instead, the level at which ribosomal proteins are downregulated correlates with the enrichment of codons that rely on s^2^U for proper decoding, indicating that these codons bias their translation in the absence of s^2^U.

It is important to note that we also saw metabolic changes in Δ*tusB* that could be contributing to the slowed growth and antibiotic tolerance phenotypes (**Figure 3**). However, if reduced metabolism were the main contributing factor, we would expect to broadly see antibiotic tolerance^44–46^. Instead, loss of s^2^U seems to induce tolerance to only antibiotics that target transcription and translation (**Figure 2**), consistent with our model that loss of s^2^U is impacting the translational capacity of the cell. It is unclear what drives these metabolic changes in Δ*tusB*, but interestingly, others have noted similar metabolic changes in response to loss of s^2^U in *Saccharomyces cerevisiae* and have postulated that 2-thiolation acts as a signal to the cell that nutrients are low ^25,29^. Loss of s^2^U thus results in a compensatory increase in synthesis of sulfur-containing amino acids^25^. Additionally, carbon metabolism is altered and shifts away from oxidative phosphorylation and towards carbohydrate storage^25^, similar to what we see in our transcriptional and proteomic analysis of Δ*tusB* (**Figure 3**).

In summary, here we identified a novel mechanism by which changes in s^2^U tRNA modification levels reduce translation of ribosomal proteins and induce slowed growth and antibiotic tolerance. Future work will be needed to identify conditions sufficient to lower tRNA modification levels, to further bolster the evidence that modulating s^2^U levels is a phenotypic response to stress that promotes bacterial survival.

## Methods and Materials

### Bacterial strains

All experiments were done with the *Yersinia pseudotuberculosis* IP2666 strain^15^. Deletion strains are generated by amplifying 1kb each of the 5’ and 3’ regions flanking the gene of interest (homology arms). The 5’ homology arm includes the start codon and 6 base pairs of the beginning of the gene and the 3’ homology arm includes 6 base pairs of the end of the gene and the stop codon. These homology arms are fused together and cloned into the pSR47s suicide vector^47^. The construct is subsequently conjugated into WT *Yersinia pseudotuberculosis* (for Δ*tusB* Δ*relA*, the *relA* pSR47s deletion construct was conjugated into the Δ*tusB Yersinia pseudotuberculosis* strain).

Sucrose selects for bacteria that have complete integration of the insert (two recombination events)^47^. Deletion of the gene was confirmed by PCR. The *tusB* rescue strain was generated by amplifying the gene and 1kb upstream and downstream of the gene from WT chromosomal DNA as a single PCR product. This was inserted into pSR47s and conjugated into the Δ*tusB Yersinia pseudotuberculosis* strain. Rescued clones were confirmed by PCR. *yrvO-mnmA* complemented and control strains were generated by transforming the plasmid constructs (pDS145 *yrvO-mnmA*, pDS220 *yrvO*(C325A)-*mnmA*, and pARA13)^30^ into Δ*tusB Yersinia* by electroporation^47^. The tRNA-Lys (UUU) complemented strains were generated by cloning the tRNA-Lys (UUU) gene into the pMB67EH plasmid downstream of a P*_tac_* promoter (inducible by IPTG treatment).

### Antibiotic susceptibility assays

To assay viability of exponential phase bacteria, overnight cultures (16h, at 26°C with rotation) were sub-cultured into 2mL LB and grown for 2h at 26°C before diluting again in 2mL LB to a starting bacterial concentration of about 10^4^ CFUs/mL. Cultures were treated with antibiotics at the MIC for bacteriostatic antibiotics (1µg/mL doxycycline, 25µg/mL chloramphenicol) or MBC_90_ concentration for bactericidal antibiotics (2µg/mL gentamicin, 2µg/mL rifampicin, or 0.1µg/mL ciprofloxacin) or left untreated. Cultures were grown for 4h at 37°C. Samples were plated for CFUs every 2h. For *yrvO-mnmA* experiments overnight cultures and subcultures were treated with 0.2% arabinose throughout the entire experiment to induce expression of the construct or as a control.

### Small RNA extraction and processing to purify tRNA

Overnight cultures (16h, at 26°C with rotation) were diluted into 50mL LB and grown at 37°C for 2h and then approximately 10^9^ CFUs were pelleted. Bacterial pellets were stored at -80°C before further processing. 1mL of Trizol was added to each cell pellet and samples were rigorously vortexed for 30 seconds, followed by room temperature incubation for 5 minutes. 200µL of chloroform was added to each sample followed by shaking by hand to mix and room temperature incubation for 3 minutes. After centrifugation at 12000xg for 15mins at 4°C, the aqueous phase was isolated and brought to a concentration of 30% ethanol to precipitate large RNAs, then applied to a spin column from the Purelink miRNA isolation kit (Thermo). After centrifugation at 12000xg for 1 minute, the flowthrough containing small RNAs was collected, brought to an ethanol concentration of 70%, and applied to a second spin column. Samples were washed twice with the kit’s wash buffer and spun dry, followed by elution in nuclease-free water. A nanodrop was used to determine RNA concentration and purity and quality was assessed by Agilent 2100 Bioanalyzer Nanochip. 1µg RNA was digested at 37°C for 6 hours in a 50µL reaction containing 5mM Tris-HCl pH 8.0, 2.5mM MgCl2, 0.1 µg/mL coformycin (adenosine deaminase inhibitor), 0.1mM deferoxamine (antioxidant), 0.1 mM BHT (antioxidant), 0.083U/µL benzonase, 0.1U/µL calf intestinal alkaline phosphatase, 0.003 U/µL phosphodiesterase I, and 50nM of 15N-dA internal standard.

### LC-MS/MS of ribonucleoside modifications

Three technical replicate injections of 200ng for each sample were analyzed by chromatography-coupled triple-quadrupole mass-spectrometry (LC-MS/MS). A Waters Acuity BEH C18 column (50 x 2.1 mm inner diameter, 1.7 micron particle size) was used with an Agilent 1290 HPLC system at 25C, 0.3mL/min flow rate, in a gradient of Buffer A (0.02% formic acid in water) and Buffer B (0.02% formic acid in 70% acetonitrile) (**see Supplemental Table 1**). The HPLC was coupled to an Agilent 6495 Triple Quad mass spectrometer with an electrospray ionization source in positive mode with 200C dry gas temperature, 11L/min gas flow, 20psi nebulizer pressure, 300C sheath gas temperature, 12L/min sheath gas temperature, 3000V capillary voltage, and 0V nozzle voltage. Product ions were detected using multiple reaction monitoring (MRM) mode, with retention times and optimal collision energies determined by synthetic standards (see **Supplemental Table 2**). The sum of 260nm UV signals of the 4 canonical nucleotides, as measured by an in-line UV detector, was used as a normalization factor.

### qPCR to measure transcriptional regulation of s^2^U modification pathway in response to antibiotics

Cultures of bacteria were grown at 37°C for 2h and treated with the MIC of antibiotics (1µg/mL doxycycline, 25µg/mL chloramphenicol, or 15µg/mL gentamicin) or left untreated. Approximately 10^7^ CFUs were pelleted and then resuspended in Buffer RLT (QIAGEN) + β-mercaptoethanol. RNA was isolated using the RNeasy kit (QIAGEN). DNA was removed using the on-column DNase digestion kit (QIAGEN). RNA was reverse transcribed using the Protoscript II First Strand cDNA synthesis kit (NEB). Approximately 30ng of cDNA was used as a template for qPCR using SYBR Green PCR Master Mix according to the manufacturer’s protocol (Applied Biosystems). Relative transcript levels were measured using the DCT and 2^-ΔCT^ method (using 16S transcript levels as an endogenous control).

### RNA-seq

Overnight cultures (16h, at 26°C with rotation) were diluted into 4mL fresh LB and grown at 37°C for 2h. Approximately 10^7^ CFUs were pelleted and then resuspended in Buffer RLT (QIAGEN) + β-mercaptoethanol. RNA was isolated using the RNeasy kit (QIAGEN). DNA was removed using the on-column DNase digestion kit (QIAGEN).

Purified RNA was shipped to Novogene (Novogene Corporation Inc), for library preparation, sequencing, and bioinformatic analysis. Sequencing reads were aligned to IP2666 *Yersinia pseudotuberculosis* genome assembly GCF_003814345.1. DESeq2 was used to determine significant differences in transcript levels between Δ*tusB* and WT samples. Genes were considered significantly differentially regulated if there was greater than 2-fold change in transcript levels and based on an adjusted p-value of .05 (p_adj_, adjusted p-value based on transcript size).

### Protein extraction

Briefly, overnight cultures (16h at 26°C with rotation) were diluted into 50mL fresh LB and then grown for 2 hours at 37C. Approximately 10^9^ CFUs were pelleted and stored in -80C. Bacterial pellets were subsequently washed in PBS then resuspended in 4X cell pellet volume of Bacterial Protein Extraction Buffer (BPER, Thermo-Fisher). After incubation for 15 minutes at room temperature, samples were centrifuged at 15,000xg for 5 mins at 4C. Supernatants (soluble fraction) and pellets (insoluble fraction) were saved at -80C. Pellets were subsequently processed to extract the insoluble fraction by washing with 1:10 diluted BPER and resuspending in 8x cell pellet volume of Inclusion Body Solubilization Reagent (Thermo-Fisher). Samples were vortexed for 1 minute and shaken on a platform shaker for 30 minutes at room temperature before being sonicated in an ice water bath for a total of 5 minutes (30 seconds on pulse, with 10 seconds off in between pulses). Samples were then centrifuged at 13,000xg for 15min at 4C and the supernatant was saved at -80°C.

### LC-MS/MS to detect relative protein expression changes in Δ*tusB* vs. WT

40µg of protein for each sample was processed for mass spectrometry using SP3 cleanup and on-bead trypsin/LysC digest to remove contaminants and digest samples into peptides. Peptides were subsequently labelled with tandem mass tags (TMT). Peptides were separated by HPLC (Thermo-Fisher Vanquish NEO UHPLC) using a self-packed fused silica column over a 90 minute gradient. Eluted peptides were sprayed into an Exploris 480 mass spectrometer. Survey scans (full ms) were acquired on an Orbi-trap within 375-1,500 m/z using a data dependent mode. The full scan MS was followed by MS/MS for the top 15 peptides in each cycle with dynamic exclusion of 45s. Precursor ions were individually isolated with 0.7 Da and fragmented (MS/MS) using an HCD activation collision energy of 36. Precursor and fragment ions were analyzed at a resolution of 120,000 and 30,000, respectively. Reporter ions at resolution of 45,000 (Turbo TMT). Raw mass spectral data files were searched using Proteome Discoverer (version 3.1, Thermo Scientific) and the GCF_003814345.1 Yersinia pseudotuberculosis IP2666pBI genome assembly. Search parameters were 10 ppm mass tolerance for precursor ions. Fixed modification was carbamidomethylation of cysteine, TMTpro on Nterm, TMTpro on lysine residues. Variable modification was oxidation on methionine residues (maximum of 3). A false discovery rate (FDR) of 1% was used to validate high confidence PSMs. Normalized protein abundances were averaged across three biological replicates. Protein abundance ratios are calculated by normalizing the average protein abundance of Δ*tusB* to the average protein abundance of WT. Pairwise comparisons were done with One-way ANOVA using a p-value of 0.05 and 2-fold change in protein abundance as cut offs for significantly differentially regulated proteins.

### Detection of misincorporation of amino acids in Δ*tusB* vs. WT

A preliminary LC/MS-MS run was done on one biological replicate using the soluble and insoluble fractions from WT and Δ*tusB* samples, similar to above (with TMT labeling). Raw mass spectral data files were searched in Proteome Discoverer v3.1 (ThermoFisher Scientific) using 3 mutations as variable modifications: Glutamate (E) changed to Aspartate (D), Glutamine (Q) changed to Histidine (H), and Lysine (K) changed to Asparagine (N). Peptides were prioritized based on relative abundance in each sample and selected in order to have a good representation of all the codons we expected there to be changes at (CAA, GAA, GAG, AAA, no mutations were identified at AAG), and codons we expected to not see increased changes at in WT compared to Δ*tusB* (CAG). A subsequent LC/MS-MS run (without TMT labeling) was done to discover additional peptides with potential misincorporations before targeting these peptides by data-independent acquisition (DIA) to determine the relative abundance of modified and unmodified peptides in each sample. 10µg of protein for each sample was processed using SP3 buffer exchange and on-bead trypsin/LysC digest. Peptides were desalted on OASIS HLB uElution plates (Waters), according to manufacturer’s instructions. Peptides were separated by a 100-minute linear gradient. MS1 scans were acquired in the Orbitrap detector of the Exploris 480 mass spectrometer from 300-1250 m/z with the following settings: RF lens setting of 50%, 120,000 resolution at 200 m/z, normalized AGC of 101%, maximum injection time set to Auto, and Easy-IC internal lockmass calibration turned on. Precursor ions in each window were fragmented by HCD at 30% NCE. DIA product ion (MS2) scans were acquired in the Orbitrap detector using the following settings: 140−1800 m/z, 30,000 resolution at 200 m/z, normalized AGC of 1000%, and a maximum injection time of 54 ms.

Raw data files were searched with Chimerys against a customized *Yersinia pseudotuberculosis* UniProt FASTA database (proteome accession UP000255087, 1263 entries), with the addition of sequences of the OmpA protein (UniProt accession P38399) containing E307D, Q309H, and Q273H substitutions. Search criteria were tryptic cleavage (maximum 2 missed), peptide length 7-30, peptide charge 2-6, 20 ppm fragment ion mass tolerance, Cys carbamidomethylation as a fixed modification, and Met oxidation as a variable modification (max 3/peptide). Peptide identifications were validated by Chimerys at 1% false discovery rate (FDR) based on an auto-concatenated decoy database search.

### Ribosome profiling

Ribosome profiling was done as described in Mohamed et al.^33^. Overnight cultures (16h, at 26°C with rotation) were diluted into 200mL LB and grown at 37°C for 5 hours before 100mL (Approximately 10^10^ CFUs) was flash frozen by spraying into liquid nitrogen. Frozen droplets of bacterial culture were stored at -80°C before 50g of frozen droplets were lysed by cryo-mill. Lysate was cleared by centrifugation at 9,000xg at 4°C for 10mins and then ribosomes were pelleted over a sucrose cushion. Approximately 12.5 AU of sample was digested with MNase to remove RNA not protected by ribosomes and then 70S monosomes were isolated by fractionation over a 10%-50% sucrose gradient. RNA (or ribosome protected fragments) was isolated by phenol chloroform extraction and then samples were run on a 15% TBE Urea gel to selectively isolate fragments within the size range of 15-45nt, which has been previously shown to be the size range of fragments protected by bacterial ribosomes^34^. cDNA libraries were prepared, sequenced and reads were mapped to the IP2666 *Yersinia pseudotuberculosis* GCF_003814345.1 genome assembly. All reads that mapped to ribosomal RNA, tRNA, and non-coding RNA were excluded. Ribosome occupancy was measured at the single codon level at the A site of the ribosome. Pause scores were calculated by taking the average ribosome density at a codon in a given gene and normalizing it to the average ribosome density across that gene for all genes in the genome.

### Codon bias analysis

Codon usage was calculated for each gene in the *Yersinia pseudotuberculosis* IP2666 genome by measuring the frequency of each codon used in each gene, otherwise known as gene codon frequency. To calculate codon Z-scores, the gene-specific codon frequency was normalized to the average global usage (all ORFs in genome) of that codon and the standard deviation of that codon. Codon Z-scores for all up and downregulated proteins (2-fold change in protein abundance and p-value <0.05), were put into a data matrix and analyzed by principal component analysis. Principal component regression analysis was done using the log2fold change in protein abundance as the dependent variable. To analyze codon usage among genes that were regulated differently at the protein level versus the transcript level, we analyzed genes that were, 2-fold upregulated at the protein level and 1-fold or lower downregulated at the transcript level, and 2-fold downregulated at the protein level and 1-fold or higher upregulated at the transcript level. All graphing and analysis was performed in GraphPad Prism.

### Codon-optimized GFP reporter

The *gfp* DNA sequence, with all CAA codons switched for CAG, was synthesized by Genewiz. The CAG optimized *gfp* was then cloned into pMMB207267 downstream of the P_tac_ (IPTG inducible) promoter and then transformed into WT and Δ*tusB Yersinia* by electroporation. Overnight cultures (16h, at 26°C with rotation) were sub-cultured into 4mL LB +1mM IPTG or untreated and grown for 8 hours at 37°C. Every 2h a 200ul sample was taken to measure optical density (OD600) and fluorescence (480/520) in a black walled 96 well plate with plate reader. GFP fluorescence is normalized to OD600 to account for bacterial density. At 4h, 1mL of sample was taken to isolate RNA to quantify *gfp* transcript levels by qPCR as described above.

### Mouse Infection

Six to eight-week-old female C57BL/6 mice were obtained from Jackson Laboratories (Bar Harbor, ME) and animal experiments were approved by Johns Hopkins Institutional Animal Care and Use Committee (Protocol #: MO19H418). Mice were inoculated with either 10^3^ (normal dose) or 10^4^ CFUs (high dose). To prepare the inoculum, bacterial cultures were grown overnight for 16-18 hours in 2X-YT (Yeast-Tryptone Broth) then diluted in sterile PBS to a final concentration of 10^4^ CFUs/mL (for normal dose) or 10^5^ CFUs/mL (for high dose). 100μl of the inoculum is injected by tail vein injection. A subset of mice is treated at 48 hours post-infection with 40mg/kg of doxycycline (720μg in 100μl sterile PBS) by intraperitoneal injection. Mouse spleens were harvested and homogenized in 1mL of sterile PBS then plated on LB agar plates to enumerate CFUs per spleen. Mice were euthanized using a lethal dose of isoflurane followed by cervical dislocation.

## Acknowledgments

We thank the Davis lab for their support and feedback on this project. We could not have done this work without the help of Bob Cole, Josh Smith, and Tatiana Boronina at the Mass spectrometry core at Johns Hopkins School of Medicine. We could also not have done the ribosome profiling without the help of Allen Buskirk, Annie Campbell, and Rachel Green at Johns Hopkins School of Medicine. We also thank Patricia Dos Santos for her advice and for generously sharing plasmids with us. We would also like to thank Bob Ernst for his help coming up with the idea for the codon-optimized reporter and Satoshi Kimura for his advice on quantifying tRNA modification levels. This work was supported by NIH (NIAID) grants AI154116 and AI175307 to K.M.D. K.L.C. was also supported by training grant AI007417-26 through NIAID.

## Author contributions

**Writing – original draft:** K.L.C; **Writing-review and editing:** K.M.D; **Conceptualization:** K.L.C, K.M.D, P.C.D, T.J.B,**; Investigation:** K.L.C, A.M; **Methodology:** T.J.B, P.C.D; **Formal Analysis:** K.L.C, T.J.B, A.M; **Funding Acquisition:** K.M.D

## Declaration of generative AI and AI-assisted technologies

The authors declare that no AI or AI-assisted technologies were used in the writing process.

## Extended Data Figures

**Extended Data Figure 1.**
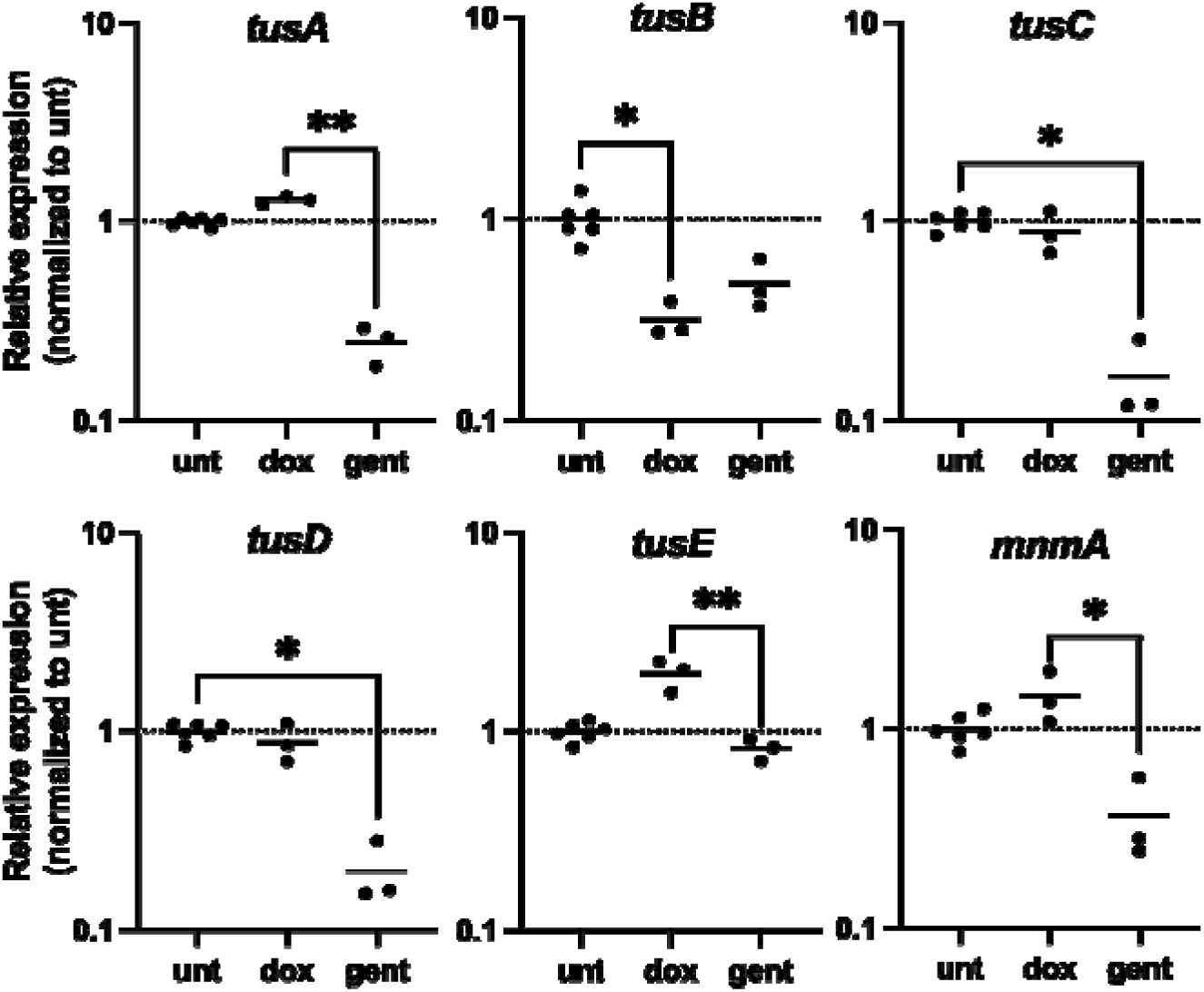
*tusB* is downregulated in response to doxycycline while multiple other genes in the 2-thiolation pathway are downregulated in response to gentamicin. Log phase bacterial cultures were grown at 37°C and treated with either 1µg/mL doxycycline, 15µg/mL gentamicin, or left untreated. After 2h RNA was isolated and relative transcript abundance of *tusA*, *tusB*, *tusC*, *tusD*, *tusE*, and *mnmA* was quantified by qPCR using 16S transcript levels as an endogenous control. Three untreated biological replicates were done in parallel with doxycycline experiments and three untreated biological replicates were done in parallel with gentamicin experiments. All six biological replicates for untreated samples and the three biological replicates for the treated samples are shown (dots). Values are normalized to the average transcript expression of the three untreated controls that were done in parallel for each experiment. Lines indicate mean relative transcript levels. Kruskal-Wallis test with Dunn’s multiple comparisons test, * p< 0.05, * p<0.01.

**Extended Data Figure 2.**
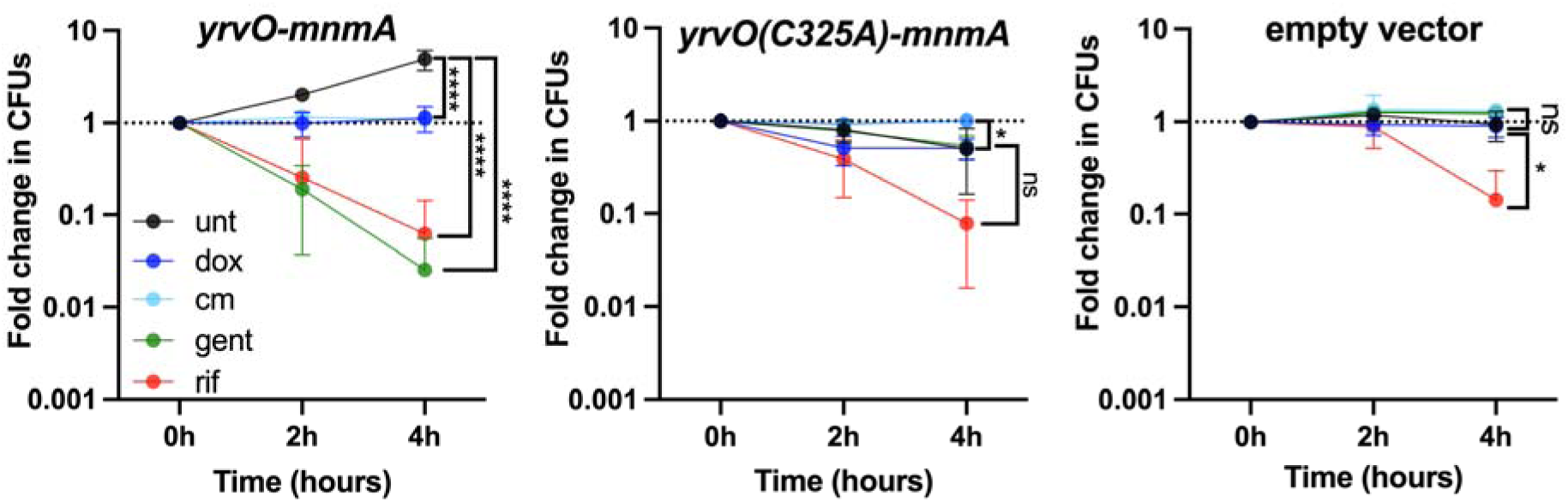
Expression of *yrvO-mnmA* restores growth and antibiotic susceptibility of. Δ***tusB.*** Log phase bacterial cultures were grown at 37°C with arabinose induction (0.2% arabinose) and sampled every 2h to enumerate CFUs (colony forming units). At the 0h timepoint, cultures were treated with either 1µg/mL doxycycline, 25µg/mL chloramphenicol, 2µg/mL gentamicin, 2µg/mL rifampicin, 0.1µg/mL ciprofloxacin, or left untreated. Fold change in CFUs are shown by normalizing each timepoint to the 0h timepoint. Two-way ANOVA with Tukey’s multiple comparisons. Asterisks denote significant p-values as follows: **** p <0.0001 and * p < 0.050. **A.** Δ*tusB* + pDS145 (pBAD plasmid containing yrvO-mnmA under the regulation of P_ara_ promoter), three biological replicates same as shown in Figure 2. **B.** Δ*tusB* + pDS220 (pBAD plasmid yrvO(C325A)-mnmA under the regulation of the P_ara_ promoter), three biological replicates. **C.** Δ*tusB* + pARA13 (pBAD empty vector control), two biological replicates.

**Extended Data Figure 3.**
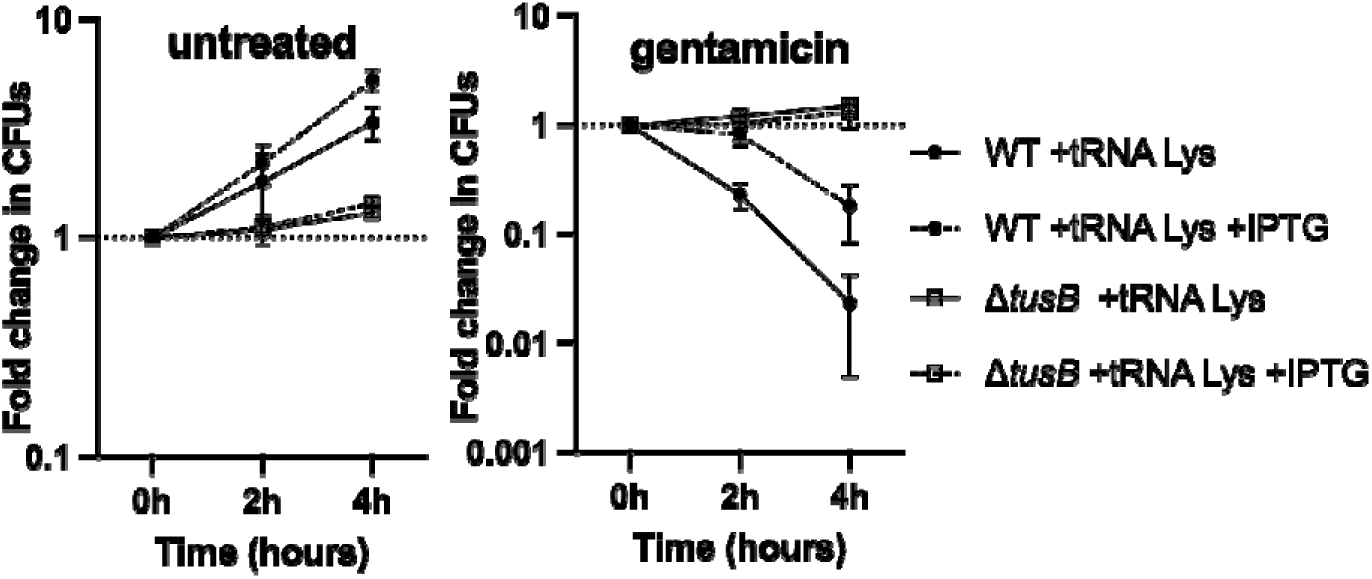
Complementing Δ*tusB* with tRNA-Lys (UUU) overexpression does not rescue slow growth or antibiotic tolerance phenotype. Log phase bacterial cultures were grown at 37°C with 1mM IPTG to induce expression of tRNA-Lys (UUU). Cultures were sampled every 2h to enumerate CFUs (colony forming units). At the 0h timepoint cultures were treated with 2µg/mL gentamicin (B) or left untreated (A). Fold change in CFUs are shown by normalizing each timepoint to the 0h timepoint.

**Extended Data Figure 4.**
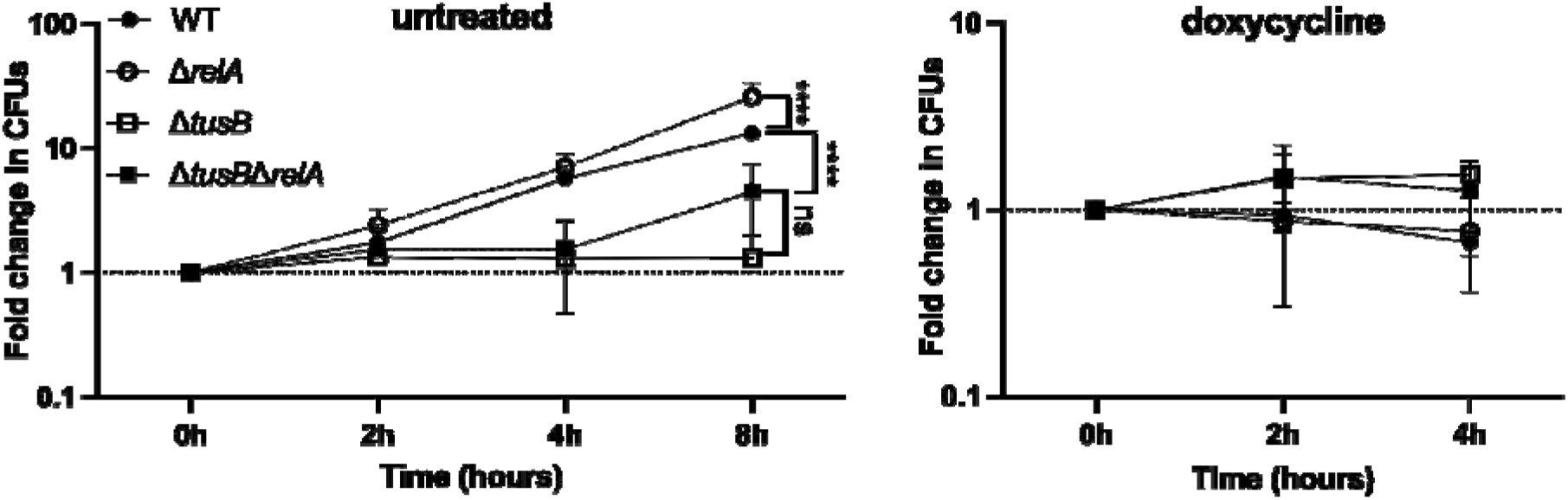
Deletion of *relA* does not restore growth or doxycycline susceptibility of Δ*tusB*. Log phase bacterial cultures were grown at 37°C and sampled every 2h to enumerate CFUs (colony forming units). At the 0h timepoint, cultures were treated with either 1µg/mL doxycycline or left untreated. Fold change in CFUs are shown by normalizing each timepoint to the 0h timepoint. Two-way ANOVA with Tukey’s multiple comparisons. Asterisks denote significant p-values as follows: **** p <0.0001. WT and Δ*tusB* doxycycline replicates are the same as shown in Figure 2.

**Extended Data Figure 5.**
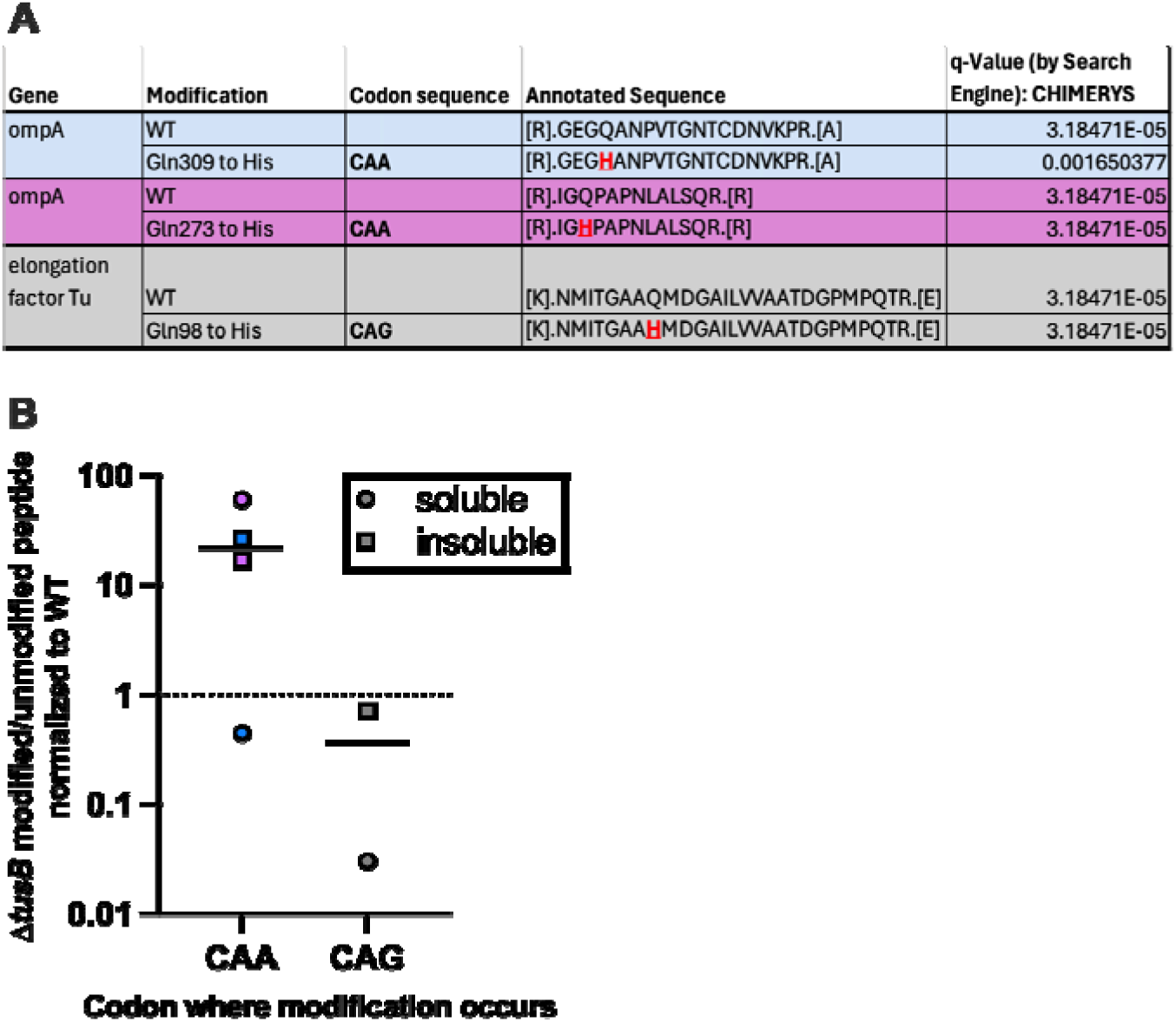
Loss of s^2^U may increase frequency of amino acid misincorporation at CAA codons. **A.** Table shows representative peptides and the modification (Glutamine Q -> Histidine H) detected in red. **B.** The abundance of each peptide was quantified by LC-MS/MS. The abundance ratio of modified to unmodified peptides in Δ*tusB* is normalized to the modified/unmodified abundance ratio in WT.

**Extended Data Figure 6.**
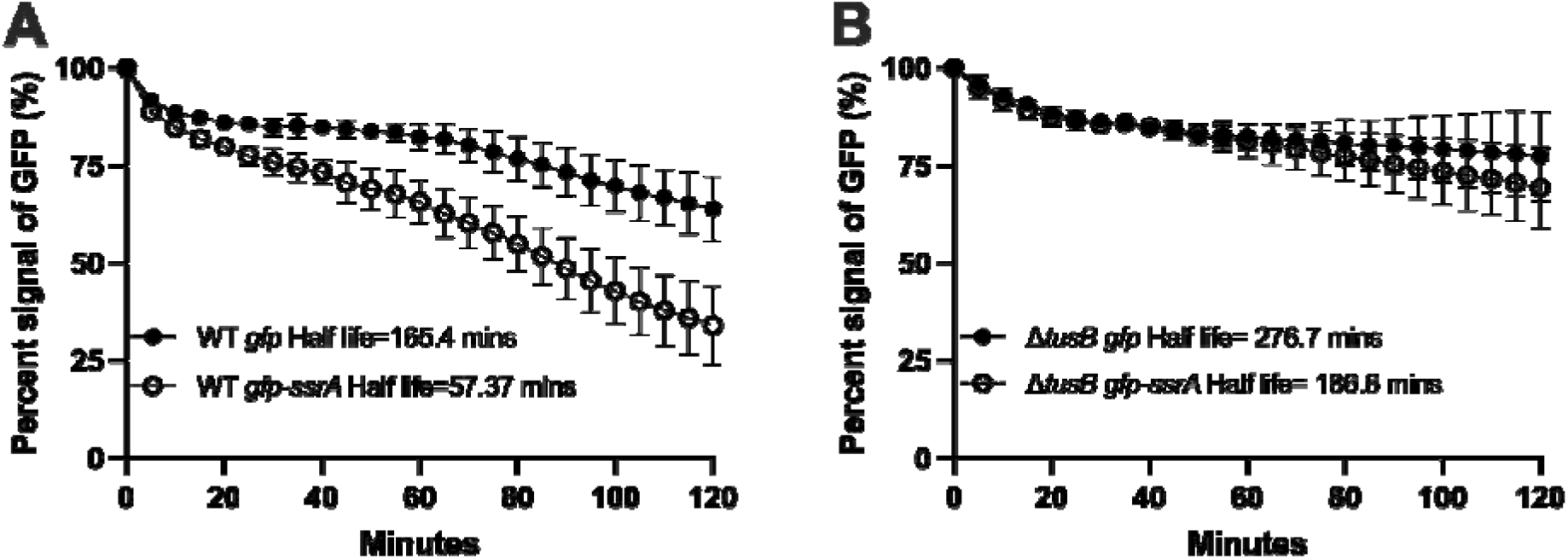
GFP protein decay is slower in *ΔtusB*. Overnight cultures were treated with 1mM IPTG to induce expression of *gfp*. After 16-18h of growth at 26°C, samples were resuspended in PBS and treated with 50 μg/mL of kanamycin (MIC) to stop translation. Fluorescence (480/520) was measured every 5 minutes for 120 minutes in a plate reader. Percent signal of the starting GFP signal (0h) is shown overtime. Half-life is calculated using the one phase decay equation using 25% GFP signal as a constraint for the plateau (limit of detection).

**Extended Data Figure 7.**
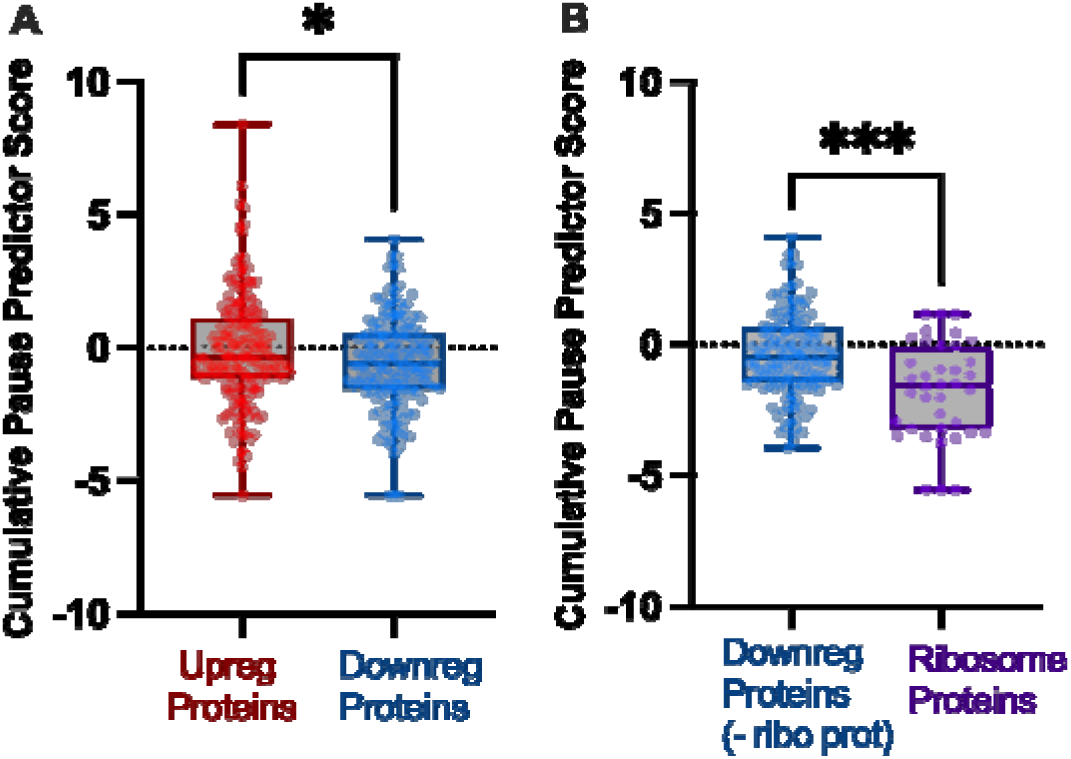
Predictor pause scores are not higher among downregulated proteins or ribosomal proteins for random codons where ribosomes do not pause at. Five codons that the ribosome does not pause more frequently at in Δ*tusB* were selected to calculate the cumulative pause predictor score as a control. Cumulative pause predictor scores were calculated by taking the sum of Ala (GCA), Arg (GAG), Asn (AAC), Cys (TGC), Gly (GGA), codon Z-scores weighted by their ribosome pause score. A. Cumulative pause predictor scores of upregulated proteins and downregulated proteins. Mann Whitney test, * p < 0.05. B. Cumulative pause predictor scores of ribosomal proteins and all other downregulated proteins *** p = 0.001

**Extended Data Figure 7.**
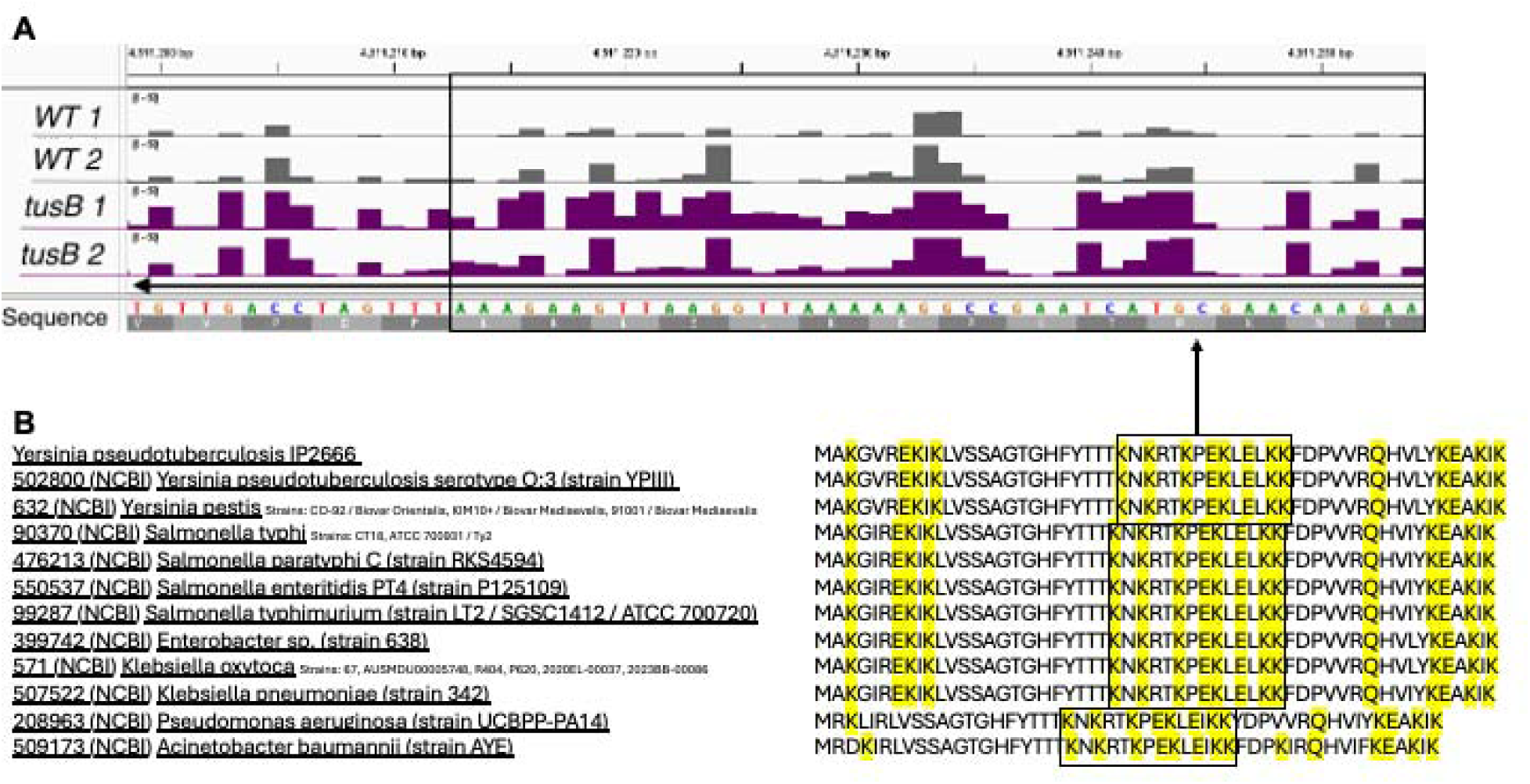
Sequences were ribosomes pause frequently at are conserved among bacterial species. **A.** Representative image of ribosome density across a sequence in the *rpmG* (ribosomal subunit L33) gene. **B.** Amino acid sequences of *rpmG* were pulled from UniProt for multiple different bacterial species. Highlighted are the residues ribosomes are predicted to pause more frequently at.

**Extended Data Table 1.** Ribosomal protein genes are downregulated at the protein level in Δ*tusB* (vs WT), but few are differentially regulated at the transcript level. All 54 ribosomal protein genes and their transcript abundance in Δ*tusB* compared to WT (as measured by RNA-seq) and their protein abundance in Δ*tusB* compared to WT (as measured by mass spectrometry). NA denotes when there was no significant difference (by p-value) detected by RNA-seq.

| Name | Gene | RNA fold change | RNA log2fold change | p<sub>adj</sub> (RNA) | Protein abundance ratio | Protein log2fold change | p-value (protein) |
|---|---|---|---|---|---|---|---|
| 30S ribosomal protein S1 | rpsA | 1.36604399 | 0.45000423 | 0.00048574 | 0.348 | -1.5228408 | 0.04431992 |
| 30S ribosomal protein S10 | rpsJ | NA | NA | NA | 0.547 | -0.8703873 | 0.00234407 |
| 30S ribosomal protein S11 | rpsK | NA | NA | NA | 0.418 | -1.2584252 | 0.00172815 |
| 30S ribosomal protein S12 | rpsL | NA | NA | NA | 0.32 | -1.6438562 | 0.02426175 |
| 30S ribosomal protein S13 | rpsM | NA | NA | NA | 0.255 | -1.9714308 | 0.0000995855 |
| 30S ribosomal protein S14 | rpsN | NA | NA | NA | 0.235 | -2.0892673 | 0.00487384 |
| 30S ribosomal protein S15 | rpsO | 0.83619814 | -0.2581282 | 0.10124687 | 0.206 | -2.2792838 | 0.00757242 |
| 30S ribosomal protein S16 | rpsP | NA | NA | NA | 0.269 | -1.8943219 | 0.01653084 |
| 30S ribosomal protein S17 | rpsQ | 1.37130023 | 0.45556044 | 0.02096267 | 0.275 | -1.8624965 | 0.00587481 |
| 30S ribosomal protein S18 | rpsR | NA | NA | NA | 0.382 | -1.3883555 | 0.11602157 |
| 30S ribosomal protein S19 | rpsS | NA | NA | NA | 0.263 | -1.9268653 | 0.00020635 |
| 30S ribosomal protein S2 | rpsB | 0.73455791 | -0.4450458 | 0.00429957 | 0.395 | -1.3400754 | 0.20138705 |
| 30S ribosomal protein S20 | rpsT | NA | NA | NA | 0.271 | -1.8836352 | 0.00042677 |
| 30S ribosomal protein S21 | rpsU | 0.68565073 | -0.5444782 | 0.00013317 | 0.196 | -2.3510744 | 0.00026588 |
| 30S ribosomal protein S3 | rpsC | NA | NA | NA | 0.371 | -1.4305089 | 0.11174822 |
| 30S ribosomal protein S4 | rpsD | NA | NA | NA | 0.463 | -1.1109159 | 0.00152108 |
| 30S ribosomal protein S5 | rpsE | NA | NA | NA | 0.387 | -1.3695945 | 0.06083988 |
| 30S ribosomal protein S6 | rpsF | NA | NA | NA | 0.761 | -0.3940316 | 0.59048579 |
| 30S ribosomal protein S7 | rpsG | NA | NA | NA | 0.338 | -1.5649048 | 0.00030091 |
| 30S ribosomal protein S8 | rpsH | NA | NA | NA | 0.466 | -1.1015981 | 0.00433927 |
| 30S ribosomal protein S9 | rpsI | NA | NA | NA | 0.332 | -1.5907449 | 0.11311814 |
| 50S ribosomal protein L1 | rplA | NA | NA | NA | 0.449 | -1.1552126 | 0.00419147 |
| 50S ribosomal protein L10 | rplJ | 0.7324748 | -0.4491423 | 0.02923479 | 0.59 | -0.7612131 | 0.07999833 |
| 50S ribosomal protein L11 | rplK | NA | NA | NA | 0.975 | -0.0365259 | 0.97643422 |
| 50S ribosomal protein L13 | rplM | 0.74307233 | -0.428427 | 0.00129496 | 0.298 | -1.7466158 | 0.00246695 |
| 50S ribosomal protein L14 | rplN | NA | NA | NA | 0.465 | -1.1046974 | 0.00269274 |
| 50S ribosomal protein L15 | rplO | NA | NA | NA | 0.388 | -1.3658714 | 0.00252536 |
| 50S ribosomal protein L16 | rplP | NA | NA | NA | 0.485 | -1.0439433 | 0.0004594 |
| 50S ribosomal protein L17 | rplQ | NA | NA | NA | 0.266 | -1.9105018 | 0.00038774 |
| 50S ribosomal protein L18 | rplR | NA | NA | NA | 0.356 | -1.4900509 | 0.0424635 |
| 50S ribosomal protein L19 | rplS | NA | NA | NA | 0.331 | -1.5950969 | 0.00154372 |
| 50S ribosomal protein L2 | rplB | NA | NA | NA | 0.398 | -1.3291597 | 0.00019869 |
| 50S ribosomal protein L20 | rplT | NA | NA | NA | 0.343 | -1.5437195 | 0.00074125 |
| 50S ribosomal protein L21 | rplU | 0.64519681 | -0.6322143 | 0.00013194 | 0.385 | -1.3770696 | 0.0066521 |
| 50S ribosomal protein L22 | rplV | NA | NA | NA | 0.347 | -1.5269924 | 0.00107999 |
| 50S ribosomal protein L23 | rplW | NA | NA | NA | 0.289 | -1.7908586 | 0.00217311 |
| 50S ribosomal protein L24 | rplX | NA | NA | NA | 0.201 | -2.3147326 | 0.00026907 |
| 50S ribosomal protein L25 | rplY | 0.57933516 | -0.7875324 | 0.00119562 | 0.66 | -0.5994621 | 0.12471253 |
| 50S ribosomal protein L27 | rpmA | 0.63606651 | -0.6528387 | 0.00336469 | 0.356 | -1.4900509 | 0.0000180606 |
| 50S ribosomal protein L28 | rpmB | NA | NA | NA | 0.281 | -1.831358 | 0.07549257 |
| 50S ribosomal protein L29 | rpmC | 1.27133672 | 0.34643557 | 0.10006984 | 0.283 | -1.821126 | 0.24651617 |
| 50S ribosomal protein L3 | rplC | NA | NA | NA | 0.25 | -2 | 0.00010761 |
| 50S ribosomal protein L30 | rpmD | NA | NA | NA | 0.275 | -1.8624965 | 0.00819852 |
| 50S ribosomal protein L31 | rpmE | NA | NA | NA | 0.518 | -0.948976 | 0.06601131 |
| 50S ribosomal protein L32 | rpmF | NA | NA | NA | 0.688 | -0.5395195 | 0.47978783 |
| 50S ribosomal protein L33 | rpmG | NA | NA | NA | 0.162 | -2.6259343 | 0.00143651 |
| 50S ribosomal protein L34 | rpmH | 0.82309168 | -0.2808809 | 0.03025432 | 0.546 | -0.8730271 | 0.44592677 |
| 50S ribosomal protein L35 | rpmI | NA | NA | NA | 0.189 | -2.4035419 | 0.00363371 |
| 50S ribosomal protein L36 | rpmJ | NA | NA | NA | 0.294 | -1.7661119 | 0.00088671 |
| 50S ribosomal protein L4 | rplD | NA | NA | NA | 0.311 | -1.6850135 | 0.00139106 |
| 50S ribosomal protein L5 | rplE | NA | NA | NA | 0.474 | -1.077041 | 0.03034906 |
| 50S ribosomal protein L6 | rplF | NA | NA | NA | 0.623 | -0.6826959 | 0.00160103 |
| 50S ribosomal protein L7/L12 | rplL | 0.834444 | -0.2610968 | 0.08072009 | 0.562 | -0.831358 | 0.50418816 |
| 50S ribosomal protein L9 | rplI | NA | NA | NA | 0.454 | -1.1392358 | 0.07865002 |

## Supplemental Information

**Supplemental Tables 1 and 2** Buffer gradient and compound table for LC-MS/MS analysis (related to Figure 1)

**Supplemental Data 1.** tRNA modification analysis (related to Figure 1)

**Supplemental Data 2.** Antibiotic susceptibility assays (related to Figure 2 and Extended Figures 2 and 3)

**Supplemental Data 3.** KEGG Analysis (related to Figure 3)

**Supplemental Data 4.** Ribosome pause scores (related to Figure 4)

**Supplemental Data 5.** CAG optimized GFP fluorescence and qPCR analysis (related to Figure 5)

**Supplemental Data 6.** RNA-seq, mass spectrometry, and codon usage analysis (related to Figures 3 and 6).

**Supplemental Data 7.** qPCR analysis for expression of s^2^U pathway genes with doxycycline and gentamicin treatment (related to Extended Data Figure 1)

**Supplemental Data 8.** Detected peptides with amino acid misincorporations (related to Extended Data Figure 4)

**Supplemental Data 9.** GFP decay in WT and Δ*tusB* (related to Extended Data Figure 5)

**Supplemental Data 10.** Mouse infection data from Figure 7.

