## Supplementary material for "Loss of the s^2^U tRNA modification induces antibiotic tolerance and is linked to changes in ribosomal protein expression": Supplementary Tables (tRNA mods).docx

**Supplementary Table 1. Buffer gradient used in the LC-MS/MS of ribonucleosides**

| **Time (min)** | **%A** | **%B** | **Flow rate (mL/min)** |
| --- | --- | --- | --- |
| 0 | 100 | 0 | 0.300 |
| 5 | 99 | 1 | 0.300 |
| 6 | 98 | 2 | 0.300 |
| 7 | 97 | 3 | 0.300 |
| 8 | 95 | 5 | 0.300 |
| 9 | 93 | 7 | 0.300 |
| 10 | 90 | 10 | 0.300 |
| 12 | 88 | 12 | 0.300 |
| 13 | 85 | 15 | 0.300 |
| 15 | 80 | 20 | 0.300 |
| 16 | 25 | 75 | 0.300 |
| 17 | 0 | 100 | 0.300 |
| 18 | 0 | 100 | 0.300 |
| 20 | 0 | 100 | 0.300 |
| 21 | 100 | 0 | 0.300 |
| 25 | 100 | 0 | 0.300 |

**Supplementary Table 2. Compound table for LC-MS/MS analysis.**

| **Compound** | **Precursor** | **Product** | **RT (min)** | **Compound** | **Precursor** | **Product** | **RT (min)** |
| --- | --- | --- | --- | --- | --- | --- | --- |
| 15N-dA | 257 | 141 | 4.6 | m5C | 258 | 126 | 1.46 |
| ac4C | 286 | 154 | 7.87 | m5s2U | 275 | 143 | 8.21 |
| acp3U | 346 | 214 | 1.37 | m5U | 259 | 127 | 4.16 |
| Am | 282 | 136 | 7.47 | m66A | 296 | 164 | 11.4 |
| Cm | 258 | 112 | 2.84 | m6A | 282 | 150 | 8.91 |
| cmnm5s2U | 348 | 141 | 2.39 | m6t6A | 427 | 295 | 10 |
| cmnm5U | 332 | 200 | 0.98 | m7G | 298 | 166 | 2.38 |
| cmo5U | 319 | 187 | 3.25 | mcmo5U | 333 | 201 | 9.3 |
| D_115 | 247 | 115 | 0.88 | mnm5s2U | 304 | 172 | 1.76 |
| Gm | 298 | 152 | 7.19 | mnm5U | 288.1 | 156.1 | 0.833 |
| ho5U | 261 | 129 | 1.18 | mo5U | 275 | 143 | 4.56 |
| I | 269 | 137 | 3.92 | ms2i6a | 382.2 | 250.1 | 17.1 |
| i6A | 336 | 204 | 16.5 | nm5s2U | 290 | 158 | 1.29 |
| io6A | 352.1 | 220 | 12.3 | preQ1 | 312 | 163 | 2.2 |
| m1A | 282 | 150 | 1.42 | Q | 410 | 163 | 5.1 |
| m1G | 298 | 166 | 7.24 | s2C | 260 | 128 | 1.93 |
| m22G | 312 | 180 | 9.71 | s2U | 261 | 129 | 4.42 |
| m2A | 282 | 150 | 6.42 | s4U | 261 | 129 | 5.033 |
| m2G | 298 | 166 | 8.0 | t6A | 413 | 281 | 12.5 |
| m3C | 258 | 126 | 1.23 | Um | 259 | 113 | 5.7 |
| m3U | 259 | 127 | 5.8 | Y | 245 | 291 | 0.9 |
