## Supplementary figures and images for "Loss of the s^2^U tRNA modification induces antibiotic tolerance and is linked to changes in ribosomal protein expression"

### SupFig1.tiff

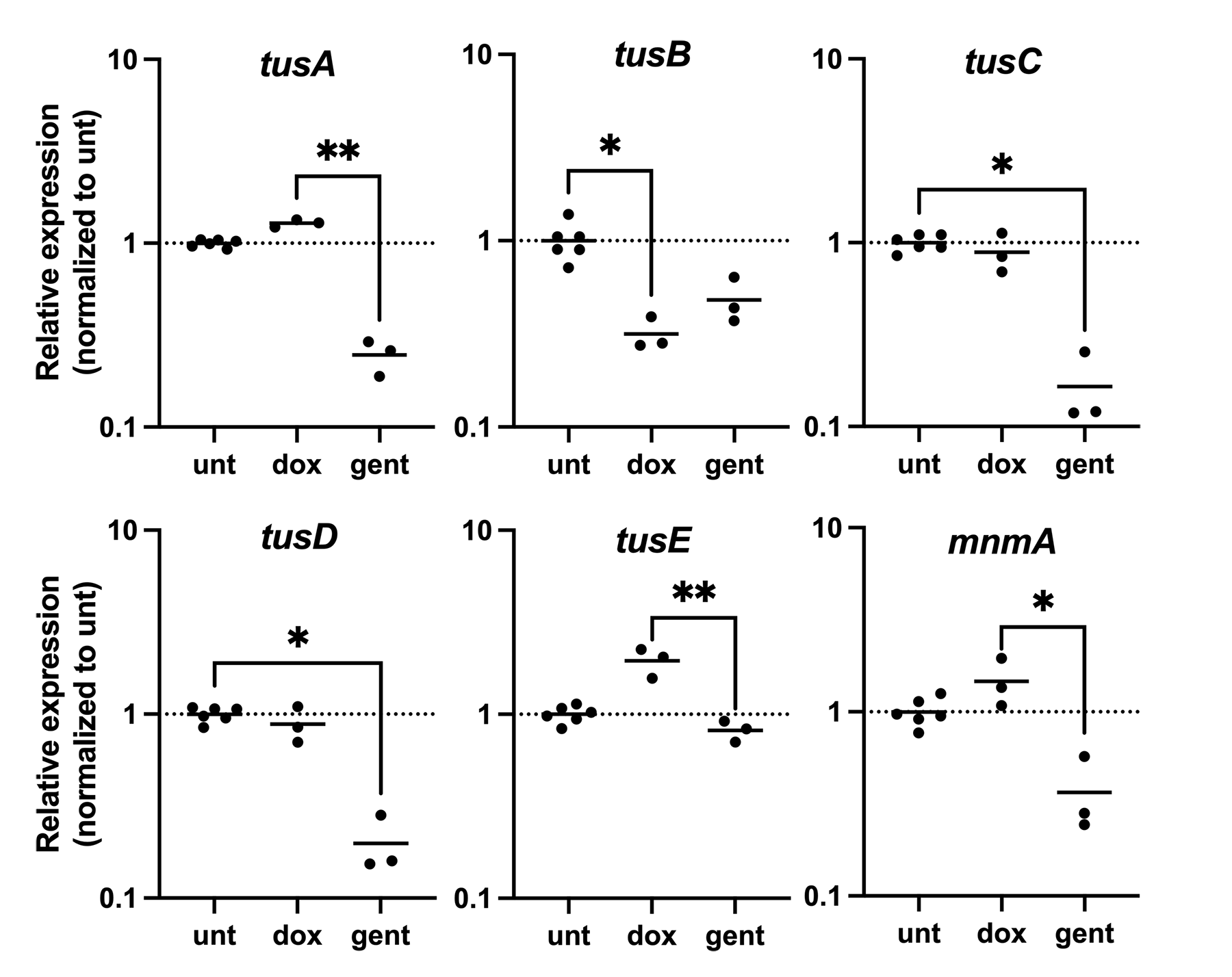

### SupFig2.tiff

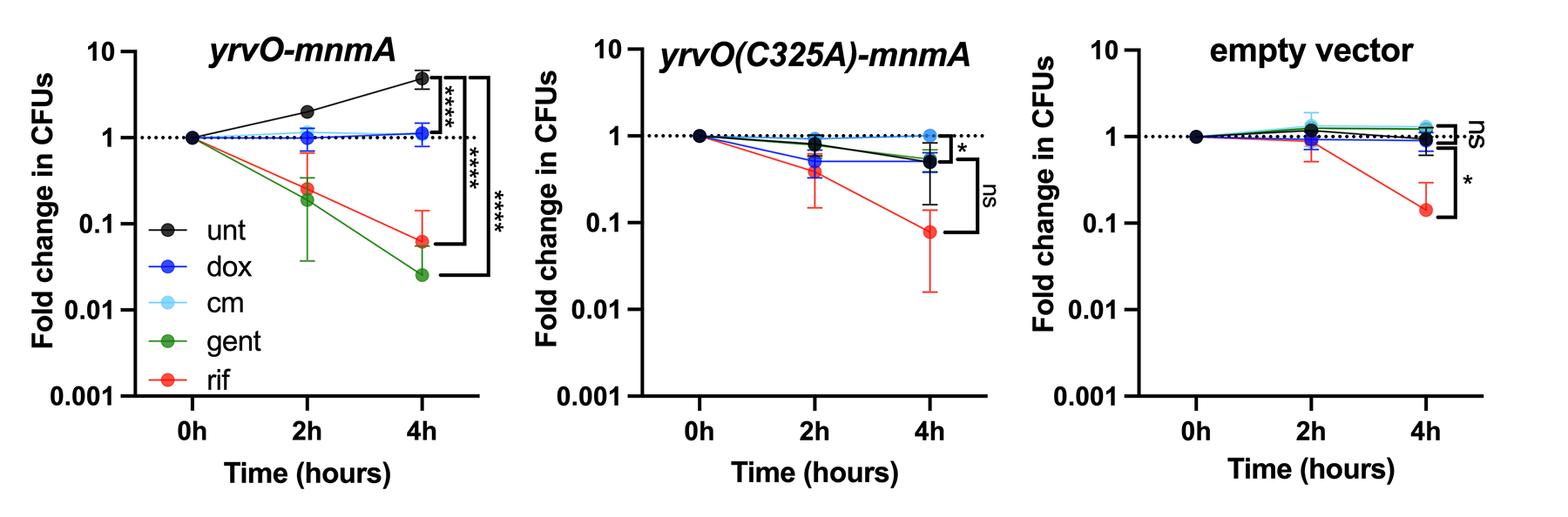

### SupFig3.tiff

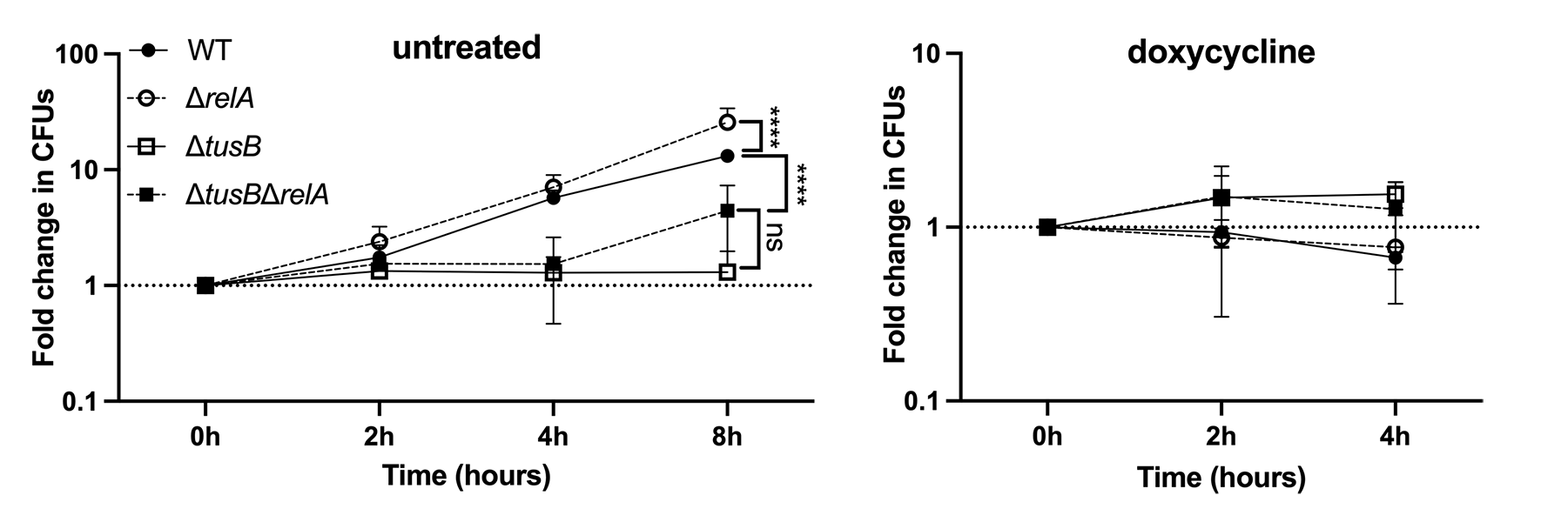

### SupFig4.tiff

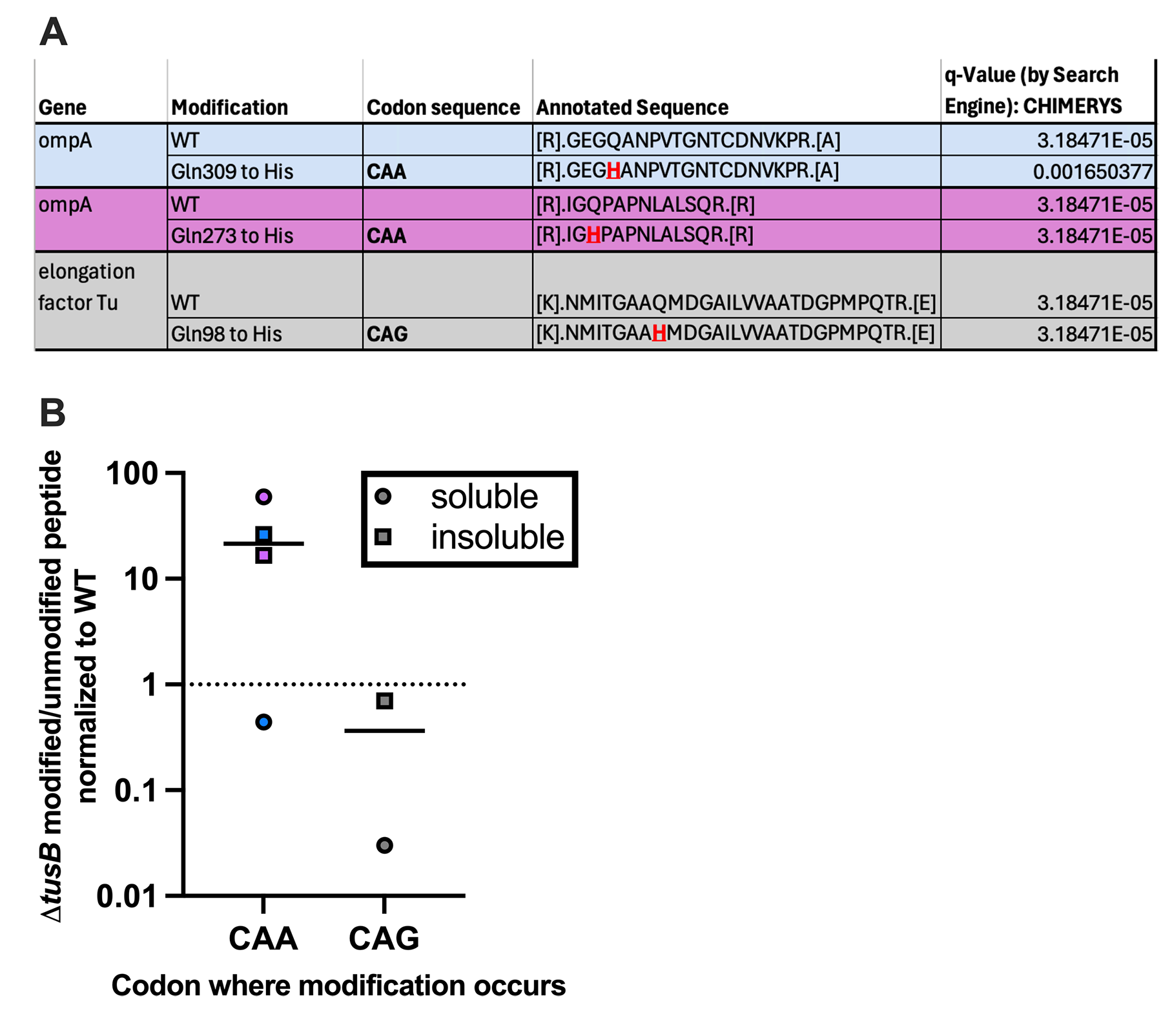

### SupFig5.tiff

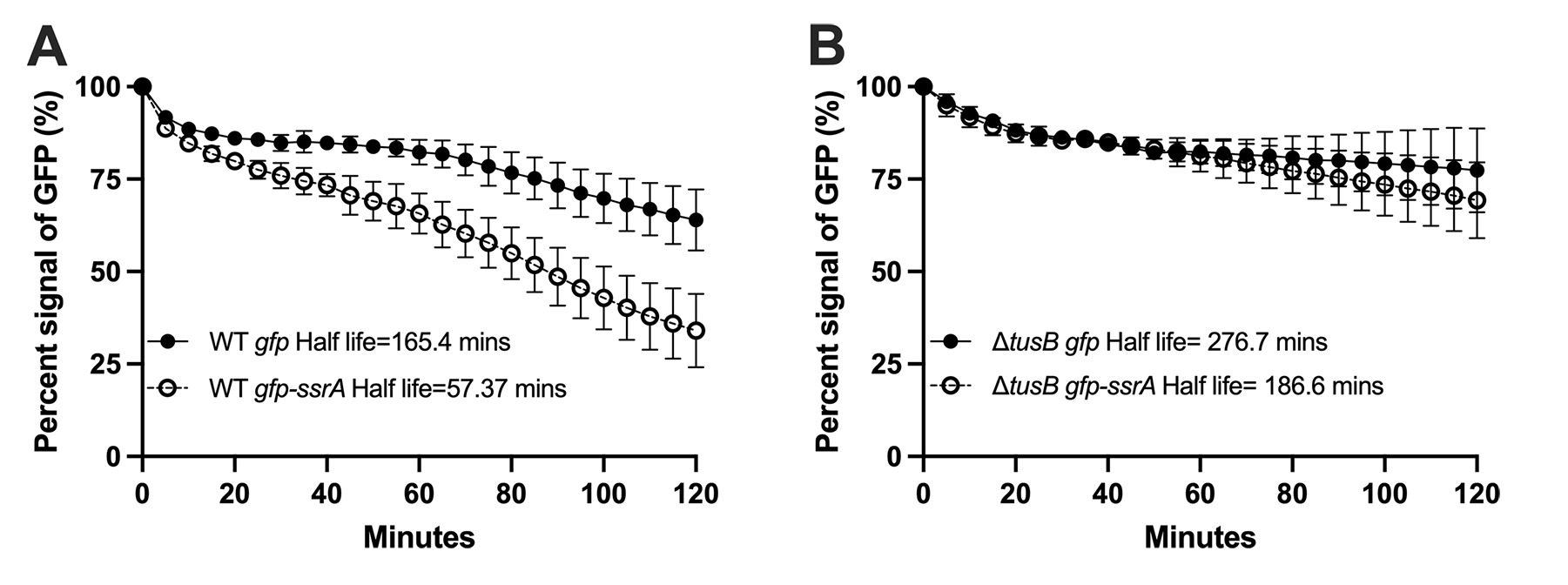

### Supp Fig 7.tif

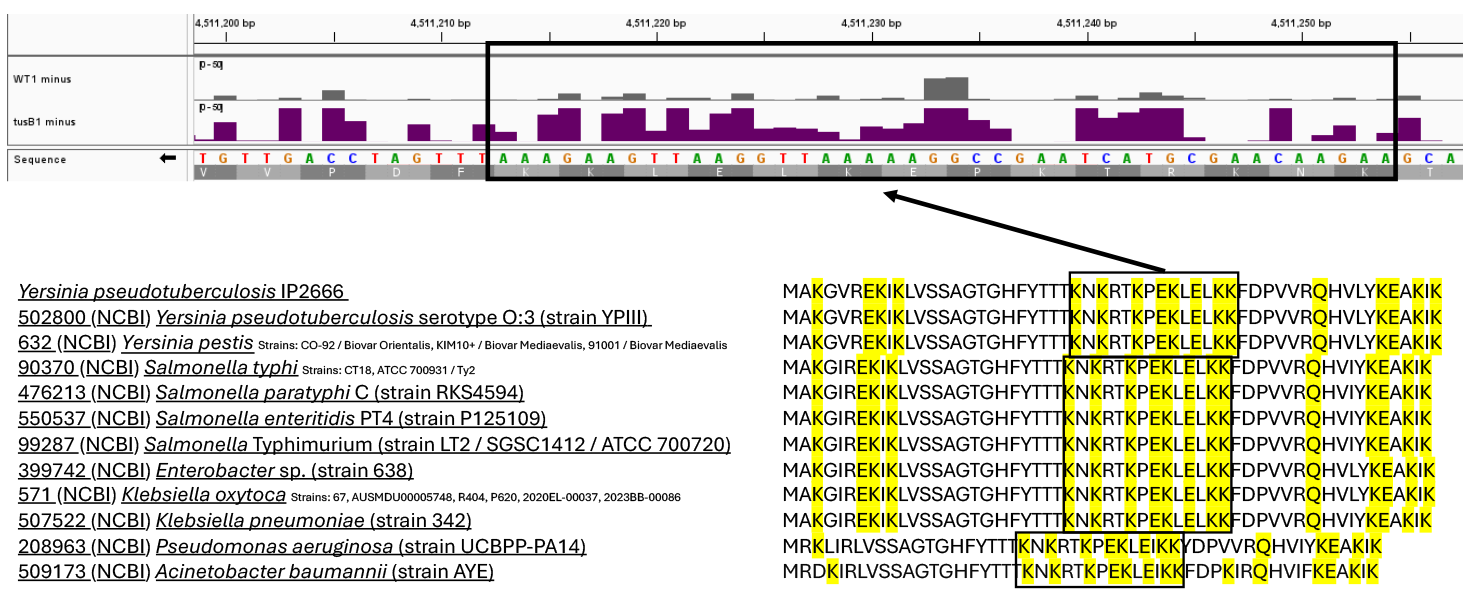

### SuppFig6.tiff

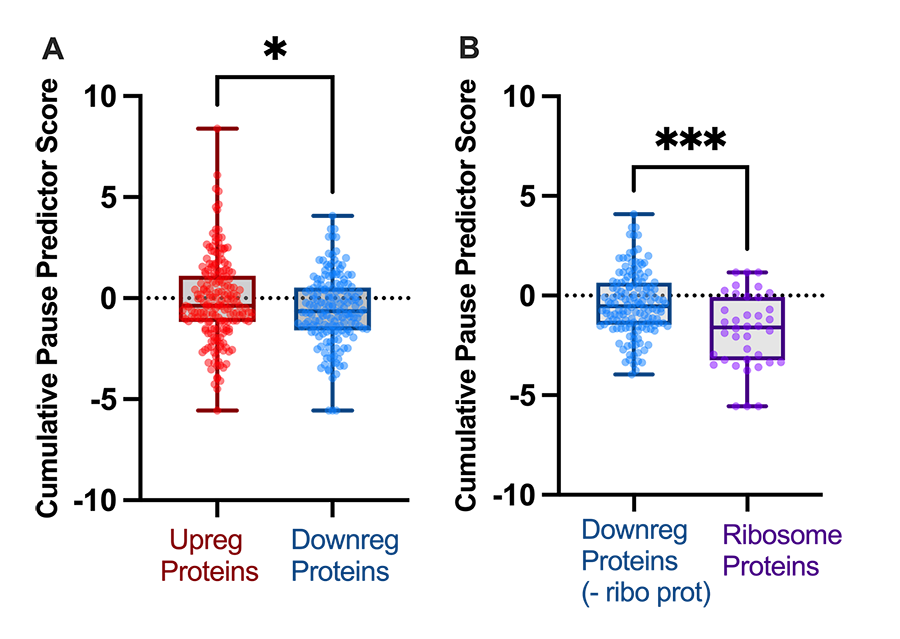
